# Single-molecule nanopore sensing of actin dynamics and drug binding

**DOI:** 10.1101/829150

**Authors:** Xiaoyi Wang, Mark D. Wilkinson, Xiaoyan Lin, Ren Ren, Keith Willison, Aleksandar P. Ivanov, Jake Baum, Joshua B. Edel

## Abstract

Actin is a key protein in the dynamic processes within the eukaryotic cell. To date, methods exploring the molecular state of actin are limited to insights gained from structural approaches, providing a snapshot of protein folding, or methods that require chemical modifications compromising actin monomer thermostability. Nanopore sensing permits label-free investigation of native proteins and is ideally suited to study proteins such as actin that require specialised buffers and cofactors. Using nanopores we determined the state of actin at the macromolecular level (filamentous or globular) and in its monomeric form bound to inhibitors. We revealed urea-dependent and voltage-dependent transitional states and observed unfolding process within which sub-populations of transient actin oligomers are visible. We detected, in real-time, drug-binding and filament-growth events at the single-molecule level. This enabled us to calculate binding stoichiometries and to propose a model for protein dynamics using unmodified, native actin molecules, demostrating the promise of nanopores sensing for in-depth understanding of protein folding landscapes and for drug discovery.

## Introduction

Actin is a ubiquitous and highly conserved ATPase found in all eukaryotes and is involved in a myriad of cellular processes including the formation of canonical eukaryotic cytoskeletal structures, cell division and cell movement^1^. Actin can also play a significant role in human diseases with rare point mutations in the actin molecule leading to aberrant aggregation and pathologies such as actin-accumulation myopathy^2^. There is also an emerging interest in the actin molecule serving as a potential drug target to stem disease. This includes targeting divergent pathogens that rely on their own actin dynamics for infectivity, such as the malaria parasite^3^, or other pathogens that utilise host cell actin for infection, such as bacteria^4^. In the human cell, actin has hundreds of binding partners, many of which facilitate the dynamic interplay between monomeric G- and filamentous F-actin forms^1^. G-actin forms a stable globular monomeric protein with intrinsic flexibility and ATPase activity^5-7^, which spontaneously forms filamentous F-actin above a critical concentration. Understanding actin dynamics, how the molecule is folded, and how it interacts with other co-factors including drugs is therefore of keen interest and advances in technologies that can explore these actin dynamics offer significant potential for the discovery of therapeutics that target host cellular processes or pathogen infection.

To date, the protein dynamics and drug binding studies of native actin have been a complex and challenging problem. For most proteins, folding is an intrinsic property of the amino acid sequence, wherein hydrophobic residues are buried to prevent aggregation, and hydrophilic residues are exposed to the aqueous environment of the cell cytoplasm. Actin does not follow this canonical folding pathway and instead relies on the species-specific chaperonin containing TCP-1 (CCT) complex. The CCT complex facilitates the addition of the co-factors ATP and divalent metal ion to fold actin into a native, polymerisation competent form^8^ without which the molecule is only partially functional^9^. Methods such as atomic force microscopy (AFM)^10^, cryo-electron microscopy (EM)^7^, crystallography^11^ and tomography^12^ have improved our understanding of actin in its G- and F-states. However, these non-time resolved approaches provide only static information and are yet to elucidate a protein folding pathway and characterise the intermediate actin folding states. While valuable information can be obtained using fluorescently labelled actin, usually by the addition of fluorophores such as N-(1-pyrene)iodoacetamide (pyrene) to the C-terminus, these chemical modifications alter the thermostability of the protein. In result, altered monomoer thermostability introduces substantial challenges in correlating polymerisation kinetics and free energy values to a native, unlabeled molecule^13, 14^. Limitations also persist when studying drug binding and its mechanisms^15, 16^. For example, the pyrenylation of actin is a common way to study drug binding, but this fluorophore only labels a minority of actin molecules and provides a readout at the level of the population, masking sub-population changes in polymerisation kinetics. Furthermore, very little information about how compounds specifically interact with actin species can be inferred from this method. Precise details of drug inhibition or stimulation and its mechanisms have traditionally relied on the structural characterisation^17, 18^, which has been instrumental in identifying the binding sites of various drugs but is time-consuming and as with other techniques provides only a static image of a protein in a non-aqueous environment.

Many of the limitations above can be addressed by using single-molecule nanopore sensing. In recent years, nanopores have been used for label-free biosensing of some of life’s fundamental building blocks such as nucleic acids^19^ and proteins^20, 21^. In a typical experiment, analytes are electrophoretically or electroosmotically translocated through a nanopore by an externally applied electric field. The translocation events lead to a characteristic temporal change of the measured ionic current passing through the nanopore. From these changes in the ionic current, one can extract information of the analyte molecular properties such as size, charge, conformational states and interactions with other biomolecules^22^. Both biological and solid-state nanopores have been used to study protein folding at the single-molecule level, revealing the conformational change and dynamics during protein unfolding^23-26^, and have also been used to observe macromolecular changes of proteins^27^. Quartz nanopipettes, a sub-class of solid-state nanopores, are low-cost and straightforward to fabricate, circumventing the technical barrier of using conventional and expensive solid-state nanopores or biological nanopores.

Here, we use quartz nanopipettes to measure real-time kinetics of the actin molecule in monomeric and polymerised state and its interaction with actin-binding drugs. Using this platform, we can distinguish between the two different macromolecular actin forms, G- and F-actin. We use the system to observe the dynamics of the unfolding of native actin using urea denaturation via measurement of thousands of single-molecule events, and we show that it is possible to observe the F-actin growth in real-time. Critically, we are able to distinguish between the binding of two drugs, Latrunculin B and Swinholide A, that prevent filament formation through different modes of action. Finally, we demonstrate the ability of nanopipette-based nanopores in drug discovery by measuring real-time drug-binding at a single molecule level, calculating the observed rate constant and the saturation concentration of Swinholide A dimer formation. Using these measurements, we propose a positively cooperative actin-binding model for the Swinholide A drug’s mode of action. The ability to perform such studies, label-free and with single-molecule resolution demonstrates the potential of using quartz nanopipettes in both the direct probing of the unfolding of complex biomolecules but also for future molecular diagnostics and drug discovery.

## Results

### Experimental configuration

Nanopipettes were fabricated using laser-assisted pulling as reported previously^28^. The pulled nanopipettes terminated with a single nanopore with an average diameter of 25 ± 4 nm (*N* = 5), as measured by Scanning Electron Microscopy (Fig. 1a and Fig. S1). Linear IV curves were obtained in the monomeric actin buffer, and the nanopore conductance was determined to be 37.0 ± 3.9 nS (Fig. S2). Given the high potassium chloride concentration (1 M), we confirmed that actin remained monomeric in the nanopore buffer (Fig. S3). In all experiments, the analyte was introduced into an external reservoir (CIS) along with a reference/ground Ag/AgCl electrode. The pipette was filled only with buffer solution, and a patch/working Ag/AgCl electrode was inserted (Fig. 1a). When an external electric field is applied, protein transport through the nanopore is a result of cooperative and competitive contributions from diffusion, electrophoretic (EP) and electroosmotic (EO) flows^29^. At pH 8.0, both the surface of the nanopipettes and actin (isoelectric point ∼5.3) are negatively charged, meaning we see no translocation events at a negative voltage (Fig. S4). We used relatively high potassium chloride concentration to suppress the EOF and to maximise the nanopore signal-to-noise ratio. At such conditions, EP is dominant and negatively charged actin molecules translocate from the bath to the inside of the nanopipette under positive bias.

**Fig. 1.**
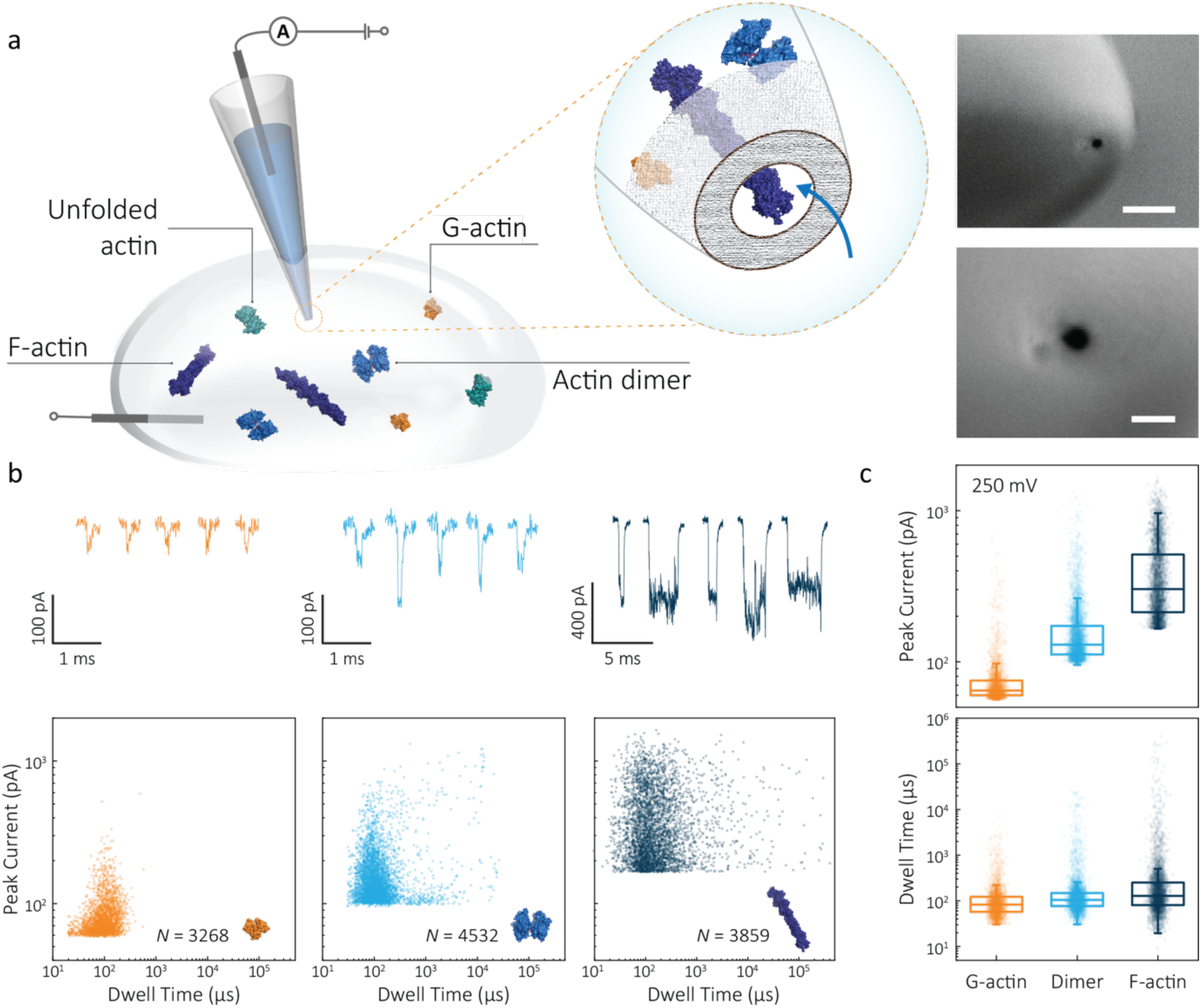
Nanopore detection of actin macromolecular states. **a** Schematic of experimental set-up of protein detection using nanopipette-based nanopore. The nanopipette and external bath were filled with a monomeric ADP-actin calcium buffer. An Ag/AgCl working electrode was inserted into the pipette, and an Ag/AgCl reference/ground electrode was fixed in the bath where the protein was introduced. Under external electric field, actin molecules in different states were translocated from the bath to the inside of the pipette through a conical nanopore at the pipette tip. Right insert: a typical SEM image showing that the nanopipette tapers down at the tip to form a nanopore with the diameter of 25 nm (scale bar 200 nm, top; 50 nm, bottom. **b** Top: nanopore translocations of three different actin species (from left to right): G-actin monomer (42 kDa, PDB: 1NWK, 800 nM), drug-induced actin dimer (PDB: 1YXQ, 500 nM) and F-actin (up to 200 actin subunits, PDB: 3G37, 800 nM). Bottom: scatterplots of current blockades vs dwell times for different actin species at 250 mV. Corresponding molecular structures are shown in the inset. **c** Box and whisker plots showing peak current and dwell time (median line and interquartile range). All data in this figure was recorded at 250 mV, sampled at 1 MHz and low-pass filtered at 50 kHz.

To demonstrate the spatial and temporal resolution of our nanopipettes, we compared the translocation characteristics of three different actin species: monomeric (G)-actin, drug-induced dimer and filamentous (F)- actin, at an applied voltage of 250 mV (Fig. 1). Two-dimensional scatterplots of dwell time (*t*_d_) vs peak current combined with the box plots (Figs. 1b and c) show we observed a cluster of events along with outliers outside the confidence interval for three species. These outliers are likely due to the transient adsorption of proteins to the nanopore surface or due to transient oligomer formation during their translocation^30^. There was an apparent expansion in dwell-time distribution when comparing G-actin to F-actin, although the mean values were close (Fig. 1c). The distribution rather than the mean value of dwell time provides key information about different features such as the molecule’s electrophoretic and electroosmotic properties or its interactions with the nanopore itself. It is noteworthy that this information relating to filament length is generally masked in ensemble averaging methods that are typically used for filament formation, such as the pyrene fluorescence assay (Fig. S3). In addition, we could also observe a distinct increase in the mean dwell time of actin dimers compared to monomers, which is likely due to a change in their respective electrophoretic mobilities.

The peak current is an essential parameter to estimate the spatial resolution of the nanopipettes, as the current blockade transients, Δ*I*_b_, are dependent on the excluded ionic volume, *Λ*, occupied by individual molecules translocated through the nanopore (eq. 1)^25^,

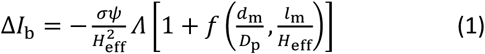

where *σ* is the solution conductivity, *ψ* is the applied voltage, *D*_p_ is the diameter and *H*_eff_ is the effective length of a nanopore, *d*_m_ is the diameter and *l*_m_ is the length of a protein molecule. The *H*_eff_ of the nanopipette was determined to be 110 ± 15 nm using 1 kb dsDNA as a standard, as shown in Fig. S9 and Method S1. *f*(*d*_m_/*D*_p_, *l*_m_/*H*_eff_) is a correction factor that primarily depends on the relative dimension between molecules themselves and the nanopore amongst others. In terms of small spherical proteins or particles, the excluded volume can be estimated as 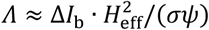 (model 1) using eq. 1. For long linear molecules *l*_m_ ≫ *H*_eff_ such as double-strand DNA (dsDNA) and polymeric or filamentous proteins, eq. 1 can be simplified as Δ*I*_*b*_ ≈ *σψ* · *A*_m_/*H*_eff_ (model 2), where *A*_m_ is the mean atomic volume per unit length of the molecule.

Model 1 was used for the analysis of the actin dimers, in which the mean peak current is 113.7 ± 15.8 pA, nearly double that of actin monomers (64.8 ± 6.7 pA). The change in current was consistent with the difference in volume between actin in these two states. For F-actin, we observed a larger mean peak current and broader distribution. This was expected since F-actin is made up of monomeric actin subunits. In the actin polymerisation process, ATP-bound monomers assemble to filaments with a 37 nm helical repeat and a 5 – 9 nm diameter^31^. Polymerisation resulted in an approximately ten-fold increase in the mean current blockade of 298.5 ± 105.8 pA when compared to actin monomers. This long-tail distribution and variety of blockade currents are likely due to the range of potential actin filament lengths possible, ranging from several subunits to micron-scale. These results demonstrate that we can use a combination of spatial and temporal analysis to reliably define the protein states (from monomer to dimer and filament) of protein-protein assemblies and their kinetics with high resolution.

### Actin unfolding using urea as a denaturant

Having established a system for studying dynamics involving the actin monomer in solution, we next sought to explore protein folding at the single-molecule level. Actin folding remains complex and poorly understood, partly due to the vast number of folding intermediates that may exist during folding pathways of any polypeptide, and also due to the intrinsic flexibility of the protein itself, which exists in a complex protein folding landscape^32^. To assess how the nanopore signal is dependent on the protein state, we performed a nanopore analysis of actin monomers exposed to 0 M to 6 M urea and bias of 250 mV. Before the measurements, the nanopipettes were tested and shown to be compatible with the chemical denaturant urea (Fig. S8), and the solution conductivity for each buffer was measured (Table S1).

The excluded volume increased with increasing urea concentration (Fig. 2). The distribution is not a standard Gaussian distribution as part of the low translocation signals were cut off by the low-pass filter, which we also see for the normalised current blockade (Δ*I*/*I*_o_) shown in Fig. S10. This trend was attributed to the increase in the exposed protein surface to the solution. Unfolding increases the solvent interaction with the protein, thus increasing the effective size of the protein and therefore contributing to a greater blockade amplitude^23^. The plot of mean excluded volume vs urea concentrations exhibits a sigmoidal shape and increases from 30.3 ± 2.5 nm^3^ to 44.0 ± 3.7 nm^3^ and corresponds to a two-state transition from folded to unfolded actin (Fig. 2b). By contrast, an opposite trend was observed for the translocation time whereby the dwell time decreases from 80.6 ± 6.4 µs to 58.1 ± 3.7 µs with increasing urea concentration (Fig. 2c) indicating the unfolded, linear actin molecules translocate faster than the folded, globular ones. We attribute this increase in translocation speed of unfolded actin to the change of charge distribution that results from the conformational changes, which in turn is associated with electrophoretic mobilities in nanopore translocation.

**Fig. 2.**
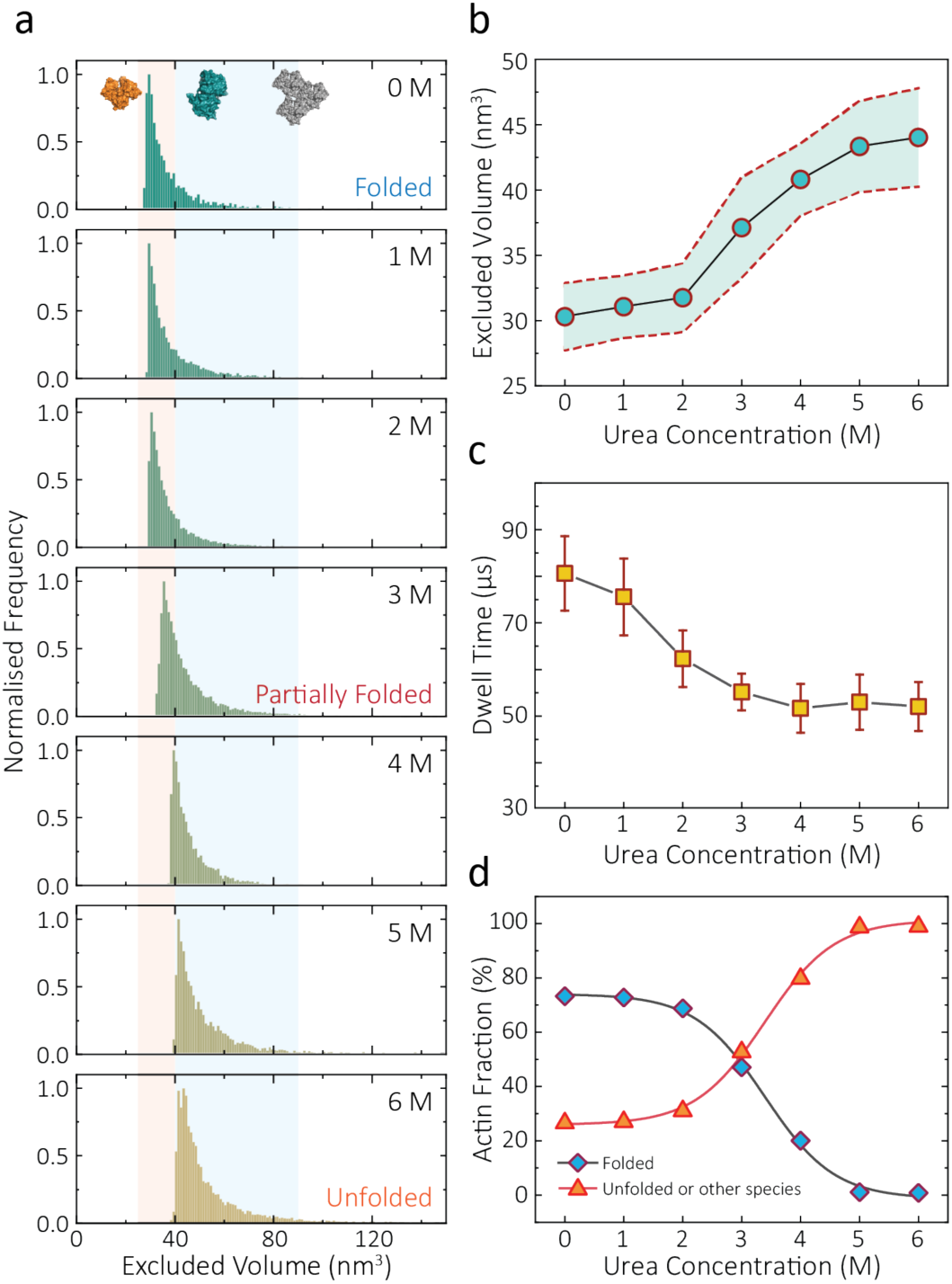
Discrimination of actin unfolding in a urea gradient using nanopipettes. **a** Normalised statistic of excluded volumes (calculated from 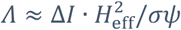) from native actin to denatured actin in increasing urea concentration. The orange boundary represents the fully folded actin state, and blue boundary determines the unfolded actin or other transient aggregates at low urea concentrations. The population shifts from a mostly native, folded state to a higher excluded volume, consistent with an unfolded state with larger hydrodynamic radius, as the urea concentration increases. Protein models for each state based on the crystal structures of G-actin (PDB: 1NWK) are shown in the inset. **b** The mean excluded volume plot exhibits a sigmoid increase when urea concentration increases, indicative of actin unfolding with an increase of hydrodynamic radius. **c** Mean dwell time decreases as the urea concentration increases. **d** Proportion of actin in different states (monomeric folded, unfolded or transient aggregated) plotted against urea concentrations shows a sigmoid curve, suggesting a two-state transition between folded (including a small fraction of aggregates) and unfolded actin.

Based on the assumption that actin is fully unfolded at 6 M urea, a threshold of 40 nm^3^ for the excluded volume at 6 M was used to define the transition between folded (< 40 nm^3^) and unfolded (> 40 nm^3^) actin. This value is around 10 nm^3^ less than reported at 0 M urea with no applied voltage as we see a slight decrease in excluded volume upon addition of voltage (Fig. S12). At 0 M urea, around a quarter (26.8 %) of translocation events have a larger amplitude than the folded monomer, which can be attributed to transient oligomers, with the majority of the population (73.2 %) forming natively folded monomers. From 0 M to 2 M urea, folded actin monomers and transient oligomers dominate, while at 3 M and above a sharp jump in the excluded volume is observed, suggesting a change in actin behaviour as a result of unfolding. As can be seen in Fig. 2d, actin unfolding is a two-state pathway in which equilibrium is reached at approximately 5 M urea, whereby we assume the majority of the population is unfolded, in agreement with previous studies using fluorescence assays^33^. Conventional analysis of protein unfolding using the denaturant exhibits a linear plot of unfolding Δ*G* vs [denaturant] in the transition zone^34^. By this linear extrapolation, we can determine the free energy of actin unfolding to be 7.74 kJ mol^−1^ at 25°C without any denaturant (Method S2, Fig. S11). It should be noted that this value is buffer- and environment-specific and is, therefore, an estimate of free energy unfolding using this nanopore platform. Despite this, these data suggest that this single-molecule statistical approach is able to describe well the dynamic changes in protein conformations measured from translocation events and can, therefore, be considered a complementary approach to traditional methods, such as fluorescence.

### Voltage-dependence on protein conformation

The applied voltage can often play a critical role in the translocation of proteins by affecting protein mobility and conformation^23, 35^. For example, the transport of proteins through the nanopore is governed by bulk diffusion, EP and EO flow and therefore the large electric field generated at the tip of the nanopipette can significantly alter the velocity and direction of the molecule and even the conformation. The effective velocity of protein transport can be described as^29^:

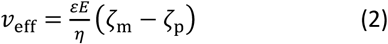

where *ε* = *ε*_0_*ε*_*r*_, *η* is the solution viscosity, *E* is the strength of the electric field, *ζ*_m_ and *ζ*_p_ is the zeta potential of the protein and walls of the nanopore, respectively. We minimised the electroosmotic component using high potassium chloride concentration, confirmed by recording translocation events only at positive bias. By recording at positive bias, only the negatively charged actin was transported from the external bath to the inside of the pipette, irrespective of buffer components (Fig. 3a). Unfolded actin translocates faster than the folded monomer within the voltage range 150 – 350 mV, independent of solution viscosity (Fig. 3b). By plotting effective velocities (*v*_eff_ = *H*_eff_/*t*_*d*_) vs applied voltages, we can extract the slope (∂*v*_eff_/ ∂*ψ*) and therefore give the zeta potential of actin using an available model (−36.0 mV for folded actin and −72.3 mV for unfolded actin, Method S3)^36^. This indicates that the unfolding process leads to a net increase in charge resulting in higher zeta potential, in agreement with the altered behaviour in native gel electrophoresis of actin^37^.

**Fig. 3.**
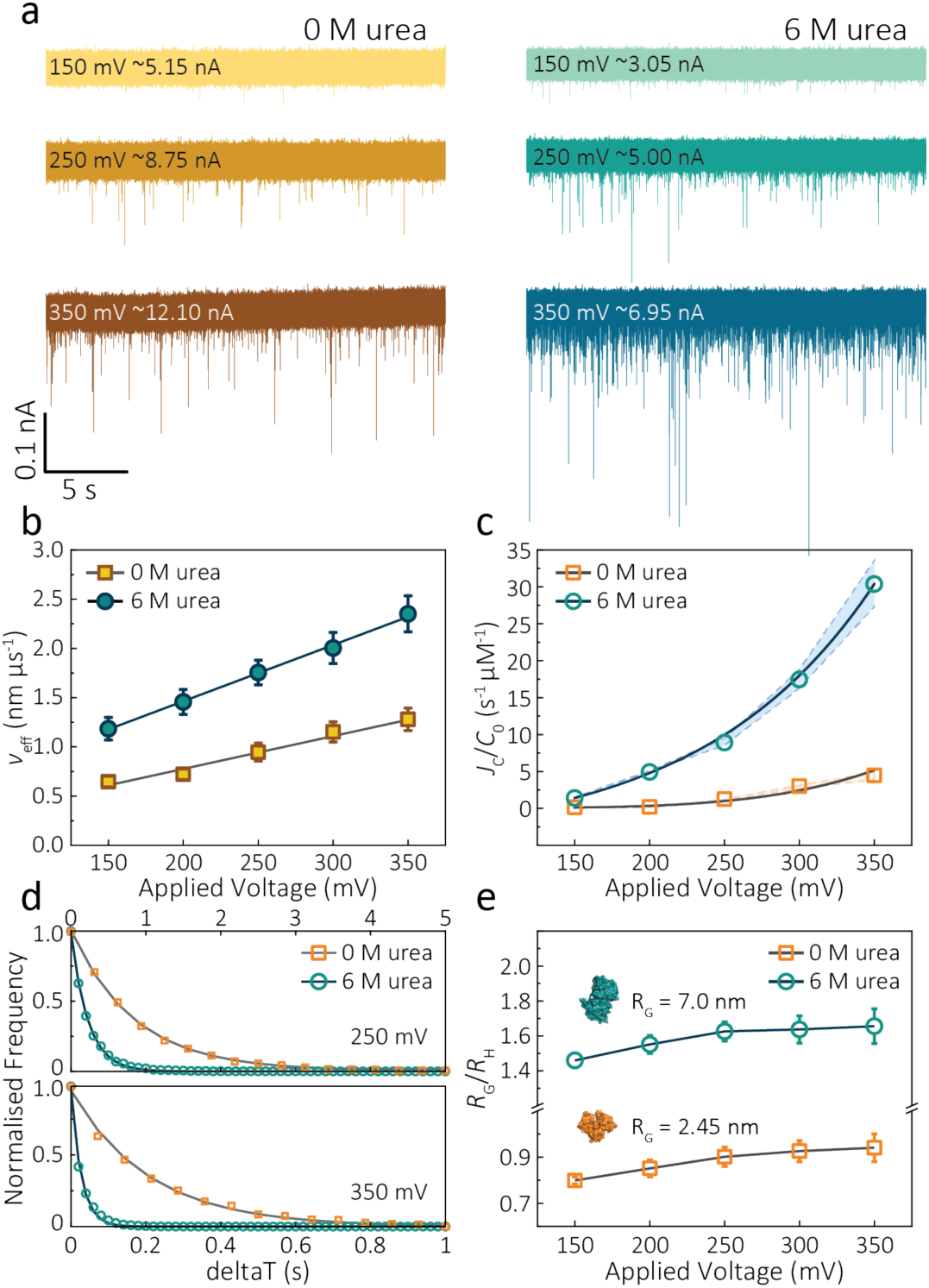
Voltage dependence upon actin translocation through nanopipettes. **a** Voltage-dependent ionic current traces show the translocation spikes of 800 nM folded actin in 0 M urea and unfolded actin in 6 M urea. Open currents (*I*_o_) for each voltage were marked upon each baseline. **b** Effective velocity (*H*_eff_/dwell time) for both folded and unfolded actin shows a linearly voltage-dependent increase when increasing the applied voltage. **c** Normalised capture rates (*J*_c_/*C*_0_) shows an exponential function of applied voltages for both folded and unfolded actin. This nonlinear increase suggests a two-stage regime in which the entropic barriers restrict successful translocations at low voltages and electrophoretic forces dominate capture behaviours at higher voltages. **d** Normalised distributions of elapsed time between successive captured events (*δt*) for actin transport in 0 M and 6 M urea buffers at 250 mV and 350 mV. Solid lines represent a single-exponential decay fit, from which the protein flux (*J*_c_) is extracted (for **c**). **e** Plots of the ratio of the radius of gyration to hydrodynamic radius (*R*_G_/*R*_H_) illustrate that the actin shape is an oblate ellipsoid in 0 M urea buffer, but a prolate ellipsoid in 6 M urea buffer during nanopore translocation. With the voltage increases, the value of *R*_G_/*R*_H_ increases both for folded and unfolded actin. This is unchanged at high voltages, indicating the protein was stretched under a high-strength localised electric field across the pipette tip.

The capture rate is another informative parameter to help facilitate our understanding of protein conformation. The protein flux (*J*_C_) is related to an entropic barrier regime. In the entropic barrier regime, the capture rate is restricted by the height of the energy barrier, with an exponential response to applied voltages. In contrast, this flux is dominated by the electric field in the drift regime, independent of the molecular diffusion and the energy barrier. The crossover point between these two regimes depends on the barrier height and molecular length itself^38^. We see a two-stage regime: an entropic barrier regime at low voltages and a drift regime at higher voltages. The steady-state flux for high voltages can be rewritten as *J*_C_/*C*_0_ = (*μψ*)/*H*_eff_ (*μ* is the electrophoretic mobility of proteins), which corresponds to the drift-dominated regime, whereby the capture rate is directly proportional to the applied voltage. At lower voltages, however, the flux is regulated by the height of the energy barrier and exhibits an exponential dependence. The normalised flux for folded and unfolded actin as a function of applied voltages is shown in Fig. 3c. Both curves display an exponential increase, indicating that the entropic barrier plays a crucial role in the actin translocation, particularly for folded actin monomers. Further, the normalised flux for unfolded actin is higher than that of folded actin which is consistent with the folded actin having a lower negative charge and hence lower zeta potential. As a result, the capture rate increases for unfolded actin, as can be seen at two different voltages (250 mV and 350 mV, Fig. 3d).

We can relate the current blockade to the hydrodynamic radius (*R*_H_) and the radius of gyration (*R*_G_) of the protein to investigate how applied voltage influences protein conformation. We assume the protein forms a hard-sphere during nanopore translocation (Methods S4^39^). Distributions of Δ*I*/*I*_o_ for folded and unfolded actin at different voltages are shown in Fig. S12, where we can observe a decrease in Δ*I*/*I*_o_ as voltage increases. The mean calculated *R*_H_ and excluded volume as a function of applied voltages are shown in Fig. S13. Importantly, the ratio of *ρ* = *R*_H_/*R*_G_ can be used to both evaluate the structural changes of the protein during translocation but more importantly relate this to the degree of asymmetry and anisotropy in the protein^39^. We can estimate the R_G_ value of the folded monomer from the crystal structure of G-actin^40^, with a value of approximately 2.45 nm, and of the unfolded protein using a model for denatured proteins, providing an estimate of around 7.0 nm^41^. We can then use these to calculate the ratio between the *R*_G_ and *R*_H_, with *ρ* of 0.775 for a globular protein with uniform density. As *ρ* increases, the protein increases to an ellipsoid. Even in harsh denaturing conditions, unfolded protein can form a native-like organised structure rather than a disordered linear conformation^42^. We calculated *ρ* values for folded actin ranging from 0.799 ± 0.015 at 150 mV to 0.942 ± 0.057 at 350 mV (Fig. 3e), suggesting the protein is approximately defined as an oblate ellipsoid, in agreement with the stretching of proteins with a dipole moment under an electric field^25^. *ρ* for unfolded actin also exhibits a similar trend increasing from 1.460 ± 0.040 to 1.658 ± 0.097. These values are more consistent with a prolate ellipsoid. Based on these observations, the localised electric field generated at the tip of nanopipettes alters the shape or conformation of translocating proteins due to their heterogeneous charge distribution. We, therefore, suggest that low voltages should be applied during nanopore experiments to understand protein-protein or protein-drug interactions. The altered behaviour at higher voltages can be used as an advantage, enabling a better understanding of some electrostatic properties such as dipole moment, net charges and molecular conductivity. Alternatively, high electric fields can be used as a method to unfold proteins without a denaturant present to complement chemical-induced unfolding.

### Actin polymerisation in real-time

Despite a plethora of actin structures, understanding the structural flexibility of actin and its polymerisation has progressed slowly^1^. Using nanopore sensing, we were able to record actin polymerisation in real-time (Fig. 4a). The ionic current of ATP-activated actin monomers was recorded over time, and translocation events with increasing peak amplitude and capture rates were observed. This noticeable increase in peak amplitudes, shown in Fig. 4b, is sensitive to molecular volumes and conformations, indicative of filament formation. We are, therefore, able to see single-molecule events and observe the distribution of dynamic interactions without averaging the population. The distribution of peak current in Fig. S14 shows time-dependent multiple populations, indicative of a concomitant increase in both the proportion and degree of ATP-actin polymerisation. Moreover, the persistence of the lowest population (105.6 ± 16.8 pA) likely represents monomeric actin, demonstrating treadmilling of the actin filament. By measuring the IV curves before and after nanopore experiments (Fig. S15), we can be certain these changes do not originate from the interaction between the analytes and the nanopore and thus are directly related to filament formation.

**Fig. 4.**
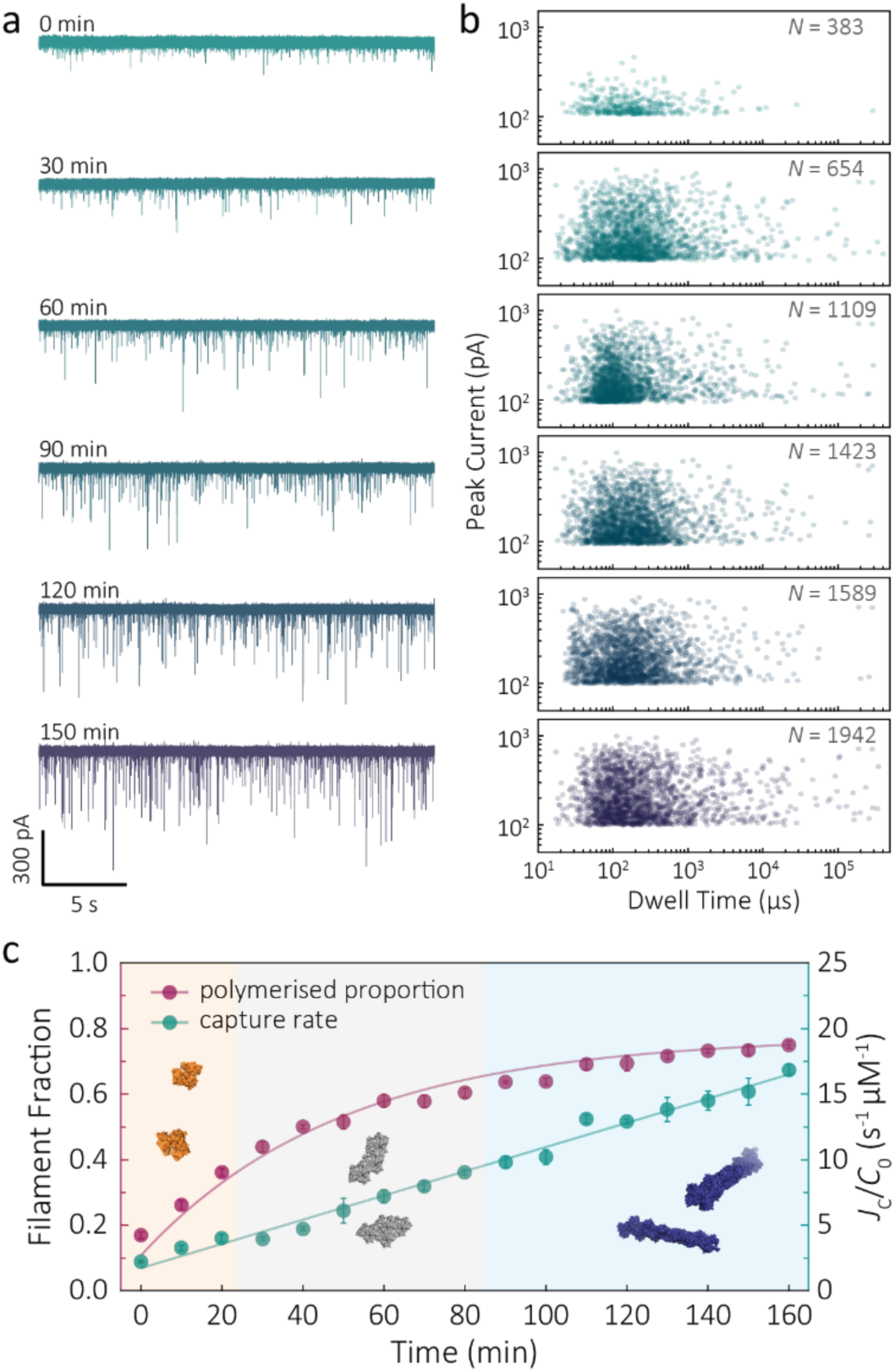
F-actin formation in real-time. **a** Time-dependent ionic current traces of actin filamentation. 1 µM actin was induced in the bath to form filaments and aggregation data was recorded in real-time. **b** Scatterplots of current blockades vs dwell times for F-actin formation at different time points corresponding to the I-t traces show in **a. c** Time-dependence of actin filament fraction (purple line) and normalised capture rate (green line) for 1 µM monomeric actin concentration. Each point was calculated every 10 min and averaged over a continuous 2 min trace. The threshold for the formation of actin oligomers and filaments was 150 pA (the mean of the second distribution in Fig. S14a). Three shadow areas and protein models (from left to right) represent actin monomers, oligomers and filaments, respectively.

Given that nanopore sensing can visually provide rich information obtained from the electro-behaviour of molecules during the transient translocation, this time-resolved method can quantitively read out the real-time properties of the analytes or their interactions. The degree of actin polymerisation can be monitored by both the filamentous fraction and event frequency (Fig. 4c). The initial fraction was around 17 % due to early actin nucleation events under high salt and ATP activation. Once polymerisation was induced, the proportion of filament increased over time and approached a maximum of 75 %, suggesting around 25 % of actin monomers are undergoing nucleotide exchange as treadmilling occurs. As expected, we see a linear increase in capture rate, a parameter which is directly related to the size, mobility and conformation of analytes. Actin filaments have higher net charge and mobility, contributing to the increasing capture events in such a drift regime. We can, therefore, use capture rate as a readout for actin polymerisation. This enables us to visualise polymerisation of native protein on a single molecule level, of critical importance to in the study of complex physiological and pathological dynamics such as protein aggregation and polymerisation.

### Actin drug binding

Given our ability to resolve unfolding actin monomers, we next sought to characterise two actin-binding drugs that have defined modes of interaction with actin as determined by X-ray crystallography. Swinholide A, a marine toxin, which binds two actin monomers in an orientation that prevents the interactions required for filament formation^44^. Latrunculins, a family of plant toxins, bind to the nucleotide-binding pocket and prevent filament formation by blocking nucleotide exchange^17, 45^.

Translocation events were recorded in the presence of either Swinholide A or Latrunculin B and compared with the monomer (Figs. 5a, S16). We observed a significant difference in the scatter plots of actin bound with Latrunculin B and Swinholide A, with much deeper current blockades and longer dwell times when binding to Swinholide A (Fig. 5b). Upon addition of excess Latrunculin B, the peak current of translocation events became more uniform at different voltages (59.1 ± 5.4 pA compared with native monomers of 64.8 ± 6.7 pA at 250 mV, Fig. S14). This suggests that incubation with excess Latrunculin B limits transient oligomeric interactions and locks actin in a monomeric state. Incubation with Swinholide A almost doubles the mean peak current from 59.1 ± 5.4 pA to 113.7 ± 15.8 pA (at 250 mV, Fig. S17) and increases the capture frequency from 1.58 ± 0.08 s^−1^ to 5.07 ± 0.16 s^−1^, whilst no observable difference can be seen between the capture frequencies of monomeric actin with or without Latrunculin B. As previously noted, this two-fold increase in current blockade is indicative of an increase in excluded volume, corresponding to the actin dimerization induced by Swinholide A.

**Fig. 5.**
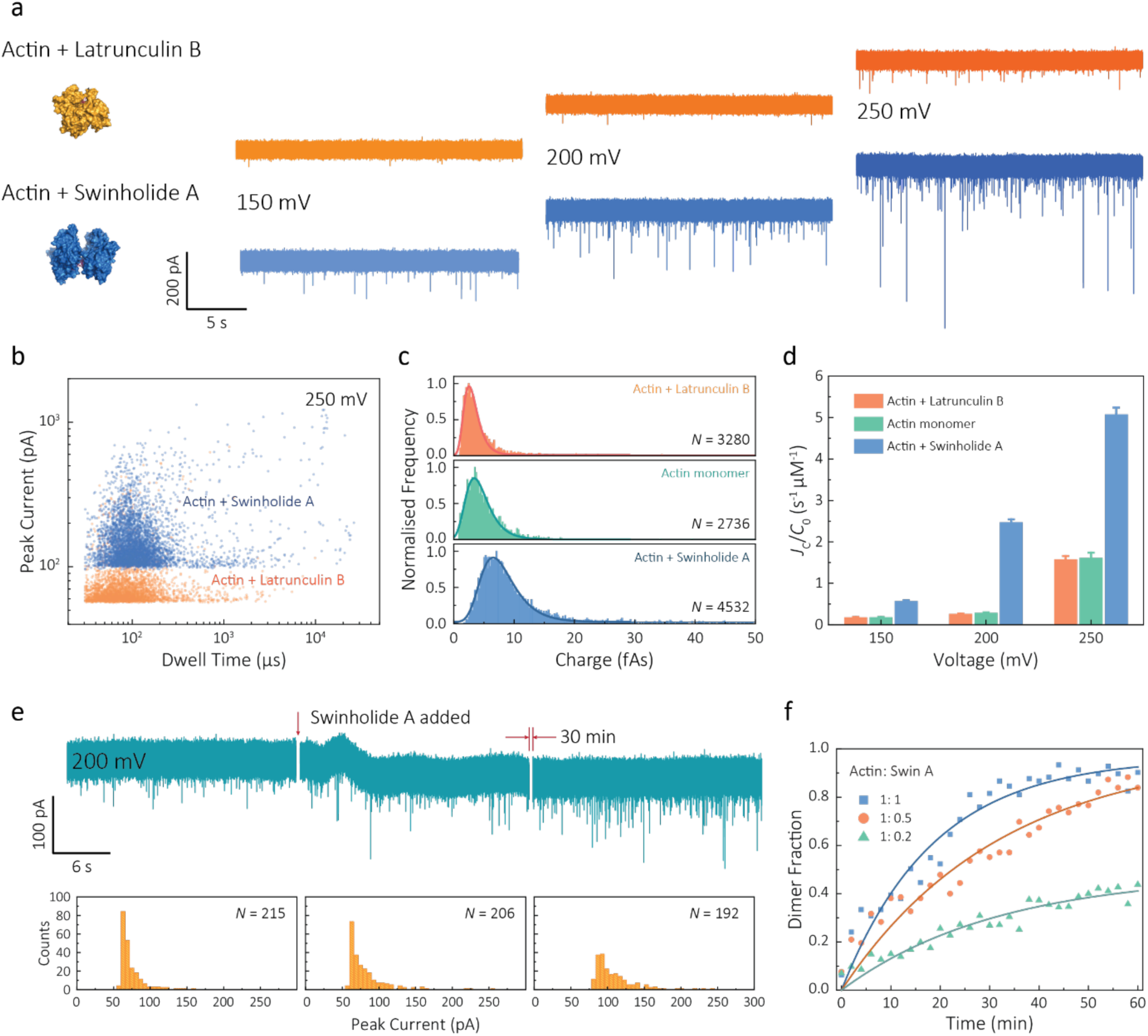
Actin drug binding assays and kinetics measurements. **a** Left: protein models based on the X-ray crystallographic structures (actin with Latrunculin B bound, PDB: 1IJJ and actin with Swinholide A bound, PDB: 10XQ). Right: typical current traces for actin bound to different filament inhibitors at 150, 200 and 250 mV. Ionic currents were recorded in monomeric buffer, 1 µM actin monomer and 10 µM filament inhibitor. **b** Scatterplots of current blockades vs dwell times for both actin monomers and dimers in the same scale at 250 mV. **c** Normalised statistic of charge drops along with a Gaussian distribution fitting for actin and actin bound with two drugs. The charge drop is equivalent to the integral area of individual translocation events. **d** Voltage dependence upon normalised capture rates of actin monomers and actin bound with two drugs. **e** Top typical current traces for drug binding kinetics collected at 200 mV. The first extract is the stable 1 µM actin monomer translocation trace. The second is the initial trace when 1 µM Swinholide A added, and the recording after 30 min is shown in the third trace. Bottom histograms represent the current blockade distributions corresponding to each trace above. **f** Time-dependent curves for 1 µM actin dimerisation with 1, 0.5, and 0.2 µM Swinholide A. The dimer fraction of each point was calculated from the statistical data in 2 min, with a threshold of 75 pA. The curves were fitted by a model of three-component interactions with positive cooperativity.

Next, we compared the charge distributions for actin alone and actin with either Latrunculin B or Swinholide A (Fig 5c). The equivalent charge, calculated by the integration of individual events, is a comprehensive parameter that includes both dwell time and peak current. As this charge profile increases, the effective charge of molecules can be considered to increase accordingly, related to the spatial excluded volume in high ionic-strength environment and molecular electrokinetics^19^. We found a charge profile of 2.5 ± 1.2 fAs for actin with Latrunculin B, 3.6 ± 2.0 fAs for monomeric actin, and 6.9 ± 2.9 fAs for actin with Swinholide A. This shift and wider distribution towards actin dimers, combined with the increase in capture frequency, is indicative of well-defined dimerisation in our measurements. The electrokinetic properties of actin monomers and dimers, including dwell time and capture rate, are shown in Fig. S17 and Fig. 5d. These properties suggest that binding with Swinholide A results in a dimeric state with a consistently higher electrophoretic mobility and net charge. This is in agreement with the crystal structure, illustrating there is a limited burial of side-chain residues and instead an overall significant increase in surface exposure to surrounding solvent molecules.

Given the noticeable differences between monomer and Swinholide A-bound actin, the drug binding efficiency of actin with Swinholide A was measured by determining the dimer fraction as a function of time (Fig. 5e). The binding was quantified at 200 mV to minimise the potential effects of changes in protein conformation caused by the electric field. Before the addition of Swinholide A, the measurement was run for at least one hour to ensure that protein flux was stable. Upon addition of two equivalents of Swinholide A, an increase occurred in the peak current distribution, as shown in Fig. 5e, consistent with the formation of actin dimers upon drug binding. The dimer fraction plateaued at around 90 % after 30 min. This can be seen in the time-dependent binding curves, Fig. 5f. Curves at different drug concentrations (with molar ratios 1:1, 1:0.5, 1:0.2, Actin: Swinholide A) were fit using a positive, cooperative model^46^. In this model, Swinholide A takes a positive effect on actin dimerisation, showing an exponential accumulation curve of the dimer fraction. The plateau can be defined as the saturated concentration of actin dimer and its corresponding rate constant can be extracted using eq. S6. Given that actin dimers have a higher flux than monomers of the same concentration (Fig. 5d), at a low drug binding ratio (e.g. 1:0.2), the calculated saturated concentration (48 %) of dimers is larger than real stoichiometric ratio (33 %). Furthermore, since the detection of dimers has a lag behind their in-situ formation due to the free diffusion and the drift by electric fields, these results reflect the apparent kinetics of protein association. This is the first time, to our knowledge, that real-time drug-binding has been visualised on a single-molecule level with native actin molecules, highlighting the potential of quartz nanopipettes for targeted drug discovery.

## Discussion

Here, we have reported on a method for single-molecule resolution analysis of a population of actin molecules that enables the analysis of changes in actin’s macromolecular or folded state as well as drug-induced structural changes. Importantly, nanopore sensing permits observations of native actin on a single molecule level without the need for chemical modifications. Other single-molecule techniques such as Atomic Force Microscopy (AFM), optical tweezers, and FRET have been used to explore protein folding accurately. However, these methods require either recombinant expression (e.g. AFM) or chemical label (e.g. optical tweezers and FRET) of the protein of interest. While suitable in some cases, for some proteins (such as actin) this is insufficient either as a result of difficulties with heterologous protein expression or interference with the protein structure or function upon addition of alternative modifications. Computational methods have, to an extent, sought to address some of these concerns, such as all-atom molecular dynamics simulations of nanopore translocations, used to explore translocations of complex native proteins^26^. Beyond computational power, any method that can proceed in the absence of modifications would be predicted to more accurately provide insights that translate from *in vitro* to *in vivo*. The combination of single-molecule data acquisition at a relatively fast rate, and use of the native protein, therefore, provides one of the key benefits of the nanopore technology over previous methods enabling close inspection of both protein folding landscapes and drug binding with enhanced statistical power.

Due to the large number of single-molecule events recorded, monitoring the behaviour of individual actin molecules in the nanopipette provides novel insight into the mechanism of urea-dependent protein unfolding. There are two current models explaining how urea unfolds proteins through the collapse of the hydrophobic core. Unfolding can be achieved either indirectly through urea interaction with water molecules or direct interactions with the protein. A number of studies have found the former to be inaccurate^47^, suggesting that a direct interaction between urea and the protein of interest is favoured. Our data support a recent model whereby, at low urea concentrations, urea binds to the protein without denaturing it, resulting in a population of dry molten globules (DMGs)^48^. We see this between 0 M and 2 M urea, where subpopulations of transient oligomers are present, and there is no observable change in the excluded volume of the protein. However, we then see substantial changes in excluded volumes between 3 M and 5 M urea that, combined with the disappearance of the subpopulations likely caused by transient interactions, agrees with the cooperative unfolding observed in previous studies^48^. For both folded and unfolded actin, the electric field applied within the nanopore can alter the protein behaviour and conformation as it translocates through the nanopore. Unlike the chemical denaturation, this conformational change is not a two-state pathway, but rather a gradual deformation or stretching the protein along the axis of the electric field^23^. Application of various electric fields has potentials to manipulate molecular conformation of macromolecules such as protein and nucleic acid in order to provide insight into the electrostatic behaviour of biological molecules.

This study provides great promise for using nanopores for drug discovery and drug binding kinetic assays. Building on previous nanopore work that has shown the ability to study the effect of drug binding on protein aggregation^49^ we applied the technology further to look at chemical and voltage induced protein unfolding and at drug binding to a monomeric species. Here, we have been able to study the effect of two different actin-binding drugs in solution and distinguish them from the monomer alone. The changes observed in charge of the actin monomer bound to Latrunculin B are distinct (Fig. 4c), despite negligible differences observed between the crystal structures of the drug-bound versus non-bound forms. The ability to resolve Swinholide A-induced changes in real-time illustrates the power of this technique for use in drug discovery. Not only can we see a distinct difference in protein behaviour, but we are also able to propose a positive cooperativity model whereby the binding of one actin molecule to Swinholide A increases its affinity to the second actin in the dimer resulting in an enhanced binding affinity. We can record these minor changes in a direct and high-throughput manner. As we look forward, efforts integrating computational analysis and classification of changes seen with a singular protein in the presence of different compounds may enable high throughput single-molecule drug screening and simultaneous prediction of the mode of interaction to be dually possible.

The sensitivity of quartz nanopipettes to both macromolecular and protein unfolding changes provide a promising outlook for studying the dynamic process of filament formation across diverse molecules using native proteins. This label-free method would work well with visualising protein aggregation, such as alpha-synuclein in real-time, which may find clinical significance in understanding the mechanism of Parkinson’s and Alzheimer’s diseases^50^. Furthermore, the high sensitivity achieved suggests the technique may provide a suitable platform not only for drug screening of specific target protein but also for a precise definition of reaction kinetics with fine granularity in solution, at a single-molecule level.

## Acknowledgements

We thank Shenglin Cai and Longhua Tang for helpful discussions in data analysis. M.D.W. is funded by the EPSRC CDT in Chemical Biology studentship. KW is funded by the EPSRC. J.B.E. is funded in part by an ERC Consolidator (NanoPD) and Proof of concept grant (Nano-PP). A.P.I. and J.B.E. are funded in part by EPSRC grant (EP/P011985/1) and BBSRC BB/R022429/1. Additional funding for the work came from the Human Frontier Science Programme (0oung Investigator Award to JB, RG00066/2016). J.B. is funded by an Investigator Award from Wellcome Trust (100993/Z/13/Z).

## Author contributions

J.B.E. and J.B. conceived and supervised the research. X.W. and M.D.W. respectively performed the nanopore experiments and prepared the protein samples and contributed equally to this work. X.L. conducted preliminary nanopore experiments. R.R. performed SEM imaging. X.W., M.D.W., J.B.E., A.P.I., R.R. and J.B. analysed the data and prepared the manuscript. All the authors discussed the results and commented on the manuscript.

## Online Methods

### Fabrication of Nanopipettes

Quartz capillaries (Intracel Ltd, UK) with an outer diameter of 1.0 mm and an inner diameter of 0.5 mm with an inner filament were cleaned using the plasma cleaner and then pulled by a laser-based pipette puller (Sutter Instrument, P-2000). The nanopipettes used in all nanopore experiments were fabricated through a two-line protocol: (1) HEAT = 775, FIL = 4, VEL = 30, DEL = 170, PUL = 80; (2) HEAT = 825, FIL = 3, VEL = 20, DEL = 145, PUL = 180. It should be noted that these parameters are instrument specific and were optimised to yield nanopore openings of 25 ± 4 nm (Fig. S1), as described in the reference.

### Preparation of actin samples

Chicken gizzard actin was purified from acetone powder using established methods. Acetone powder was suspended in 20 ml Calcium buffer G (CaBG: 2 mM Tris pH 8, 0.2 mM ATP, 0.5 mM DTT, 0.1 mM CaCl_2_, 1 mM NaAzide) per gram of powder and stirred for 30 mins at 4°C. The suspension was centrifuged at 16 000 RPM for 30 mins at 4°C in a JA25.5 rotor. The supernatant was collected, and the pellet was resuspended in the initial volume of CaBG and stirred for another 30 mins at 4°C followed by centrifugation as before. The supernatants were combined and adjusted to 50 mM KCl and 2 mM MgCl_2_ and stirred at 4°C for 1 hour to polymerise the actin. The solution was subsequently adjusted to 0.8 M KCl and stirred for a further 30 mins at 4°C before being centrifuged at 38000 RPM in an MLA 80 rotor (Beckman) for 2 hrs at 4°C to pellet the filaments. The supernatant was discarded, and the pellet was resuspended in 3-5 ml CaBG and homogenised using a 1 ml Tight Dounce Tissue Grinder (Wheaton) and dialysed into fresh CaBG overnight at 4°C. The suspension was removed from dialysis and passaged through a 26-gauge syringe 20-30 times to break down the filaments and returned to dialysis for a maximum of 3 days. The solution was centrifuged at 38000 RPM in an MLA 80 rotor (Beckman) for 2 hrs at 4°C to pellet contaminating filaments and aggregates, leaving monomeric actin in the supernatant. The supernatant was loaded onto an S200 16/600 (GE Healthcare) equilibrated with CaBG and fractions containing pure monomeric actin were analysed by SDS-PAGE, pooled and stored at 4°C for a maximum for 3 months. The sample was dialysed into fresh CaBG and centrifuged at 38000 RPM in an MLA 80 rotor (Beckman) for 2 hrs at 4°C to pellet any contaminating filaments immediately before conducting experiments.

Purified chicken actin was dialysed and diluted to 300 – 1000 nM into the nanopipette buffer (20 mM Tris pH 8, 100 mM KCl, 0.4 mM ADP, 0.1 mM CaCl_2_, 0.01% DMSO and 10 % glycerol). Actin for the urea unfolding experiments was dialysed into the nanopipette buffer supplemented with 1 – 6 M Urea. For comparison to the filament, F-actin was prepared as described ^51^. Briefly, 2.5 µM G-actin was incubated 10X Mg-EGTA exchange buffer (ME) to make Mg-ATP-Actin, which was followed by the addition of the polymerisation buffer 10X KMEI (0.5 M KCl, 0.1 M imidazole, 0.01 M EGTA pH 8.0, 10 µM MgCl_2_) before being incubated for 2 hrs at room temperature. This was then added to the nanopipette tip using wide orifice pipette tips. For observing actin polymerisation, actin was dialysed into nanopipette buffer containing 0.4 mM ATP. The polymerisation was induced by adding 10 X KME buffer (0.5 M KCl, 10 mM EGTA pH 8.0, 10 mM MgCl_2_). The preparation of the actin-LatB or actin-SwinA complex required incubation and equilibration with 10 µM LatB or 10 µM Swinholide A (dissolved in DMSO) for 30 mins before voltage application.

### Nanopore measurements and data processing

Before performing translocation experiments, 1 M KCl buffer was added into the nanopipette using a microfilm needle (MF34G, World Precision Instruments, UK). Electrodes (Ag/AgCl) were inserted into the nanopipette (working electrode) and the external bath (reference/ground electrode), respectively. Voltages from 150 mV to 350 mV were applied, and the ionic current was recorded as a function of time by Chimera amplifier VC100 (Chimera Instruments) with a sampling rate of 4.17 MHz. and a low-pass kHz digital Bessel filter of 50 kHz and analysed using custom-written MATLAB code by Edel Group (Fig. S5 and S6) unless otherwise stated. Power spectral density (PSD) plots for this low-noise configuration are shown in Fig. S7, at different applied voltages and low-pass filters. The data was filtered with a 10 kHz-100 kHz digital Bessel filter and analysed using custom-written MATLAB code by Edel Group. Protein fluxes (*J*_*c*_) were extracted from a single-exponential decay fitting of the distribution of interval time between two successive translocation events (*δt*) as previously reported^52^. All data collected at different voltages were obtained from the same nanopipette and error bars represent one standard deviation of at least three independent experiments with different nanopipettes unless stated otherwise. Traces shown were collected at 1 MHz and low-pass filtered to 50 kHz.

## References

1. Dominguez, R. & Holmes, K.C. Actin structure and function. Ann. Rev. Biophys. 40, 169–186 (2011).

2. Agrawal, P.B. et al. Heterogeneity of nemaline myopathy cases with skeletal muscle alpha-actin gene mutations. Ann. Neurol. 56, 86–96 (2004).

3. Johnson, S. et al. Truncated Latrunculins as Actin Inhibitors Targeting Plasmodium falciparum Motility and Host Cell Invasion. J. Med. Chem. 59, 10994–11005 (2016).

4. Cossart, P. & Sansonetti, P.J. Bacterial invasion: the paradigms of enteroinvasive pathogens. Science 304, 242–248 (2004).

5. Galkin, V.E., Orlova, A., Lukoyanova, N., Wriggers, W. & Egelman, E.H. Actin depolymerizing factor stabilizes an existing state of F-actin and can change the tilt of F-actin subunits. J. Cell Biol. 153, 75–86 (2001).

6. Galkin, V.E., VanLoock, M.S., Orlova, A. & Egelman, E.H. A new internal mode in F-actin helps explain the remarkable evolutionary conservation of actin’s sequence and structure. Curr. Biol. 12, 570–575 (2002).

7. Schmid, M.F., Sherman, M.B., Matsudaira, P. & Chiu, W. Structure of the acrosomal bundle. Nature 431, 104–107 (2004).

8. Stuart, S.F., Leatherbarrow, R.J. & Willison, K.R. A two-step mechanism for the folding of actin by the yeast cytosolic chaperonin. J. Biol. Chem. 286, 178–184 (2011).

9. Willison, K.R. The structure and evolution of eukaryotic chaperonin-containing TCP-1 and its mechanism that folds actin into a protein spring. Biochem. J. 475, 3009–3034 (2018).

10. Matusovsky, O.S., Mansson, A., Persson, M., Cheng, Y.-S. & Rassier, D.E. High-speed AFM reveals subsecond dynamics of cardiac thin filaments upon Ca^2+^activation and heavy meromyosin binding. Proc. Natl. Acad. Sci. USA 116, 16384 (2019).

11. Kabsch, W., Mannherz, H.G., Suck, D., Pai, E.F. & Holmes, K.C. Atomic structure of the actin:DNase I complex. Nature 347, 37–44 (1990).

12. Urban, E., Jacob, S., Nemethova, M., Resch, G.P. & Small, J.V. Electron tomography reveals unbranched networks of actin filaments in lamellipodia. Nat. Cell Biol. 12, 429–435 (2010).

13. Ooi, A., Yano, F. & Okagaki, T. Thermal stability of carp G-actin monitored by loss of polymerization activity using an extrinsic fluorescent probe. Fish. Sci. 74, 193–199 (2008).

14. Kozuka, J., Yokota, H., Arai, Y., Ishii, Y. & Yanagida, T. Dynamic polymorphism of single actin molecules in the actin filament. Nat. Chem. Biol. 2, 83–86 (2006).

15. Falsey, R.R. et al. Actin microfilament aggregation induced by withaferin A is mediated by annexin II. Nat. Chem. Biol. 2, 33–38 (2006).

16. Ueoka, R. et al. Metabolic and evolutionary origin of actin-binding polyketides from diverse organisms. Nat. Chem. Biol. 11, 705–712 (2015).

17. Morton, W.M., Ayscough, K.R. & McLaughlin, P.J. Latrunculin alters the actin-monomer subunit interface to prevent polymerization. Nat. Cell Biol. 2, 376–378 (2000).

18. Bubb, M.R., Spector, I., Bershadsky, A.D. & Korn, E.D. Swinholide A is a microfilament disrupting marine toxin that stabilizes actin dimers and severs actin filaments. J. Biol. Chem. 270, 3463–3466 (1995).

19. Ivanov, A.P. et al. On-demand delivery of single DNA molecules using nanopipets. ACS Nano 9, 3587–3595 (2015).

20. Houghtaling, J. et al. Estimation of Shape, Volume, and Dipole Moment of Individual Proteins Freely Transiting a Synthetic Nanopore. ACS Nano 13, 5231–5242 (2019).

21. Ren, R. et al. Nanopore extended field-effect transistor for selective single-molecule biosensing. Nat. Commun. 8, 586 (2017).

22. Miles, B.N. et al. Single molecule sensing with solid-state nanopores: novel materials, methods, and applications. Chem. Soc. Rev. 42, 15–28 (2013).

23. Freedman, K.J., Haq, S.R., Edel, J.B., Jemth, P. & Kim, M.J. Single molecule unfolding and stretching of protein domains inside a solid-state nanopore by electric field. Sci. Rep. 3, 1638 (2013).

24. Oukhaled, A. et al. Dynamics of completely unfolded and native proteins through solid-state nanopores as a function of electric driving force. ACS Nano 5, 3628–3638 (2011).

25. Talaga, D.S. & Li, J. Single-molecule protein unfolding in solid state nanopores. J. Am. Chem. Soc. 131, 9287–9297 (2009).

26. Si, W. & Aksimentiev, A. Nanopore Sensing of Protein Folding. ACS Nano 11, 7091–7100 (2017).

27. Yusko, E.C. et al. Single-particle characterization of Abeta oligomers in solution. ACS Nano 6, 5909–5919 (2012).

28. Cai, S., Sze, J.Y.Y., Ivanov, A.P. & Edel, J.B. Small molecule electro-optical binding assay using nanopores. Nat. Commun. 10, 1797 (2019).

29. Firnkes, M., Pedone, D., Knezevic, J., Doblinger, M. & Rant, U. Electrically facilitated translocations of proteins through silicon nitride nanopores: conjoint and competitive action of diffusion, electrophoresis, and electroosmosis. Nano Lett. 10, 2162–2167 (2010).

30. Li, W. et al. Single protein molecule detection by glass nanopores. ACS Nano 7, 4129–4134 (2013).

31. Egelman, E.H. A tale of two polymers: new insights into helical filaments. Nat. Rev. Mol. Cell Biol. 4, 621–630 (2003).

32. Kuznetsova, I.M., Povarova, O.I., Uversky, V.N. & Turoverov, K.K. Native globular actin has a thermodynamically unstable quasi-stationary structure with elements of intrinsic disorder. FEBS J. 283, 438–445 (2016).

33. Levitsky, D.I., Pivovarova, A.V., Mikhailova, V.V. & Nikolaeva, O.P. Thermal unfolding and aggregation of actin. FEBS J. 275, 4280–4295 (2008).

34. Santoro, M.M. & Bolen, D.W. A test of the linear extrapolation of unfolding free energy changes over an extended denaturant concentration range. Biochemistry 31, 4901–4907 (1992).

35. Freedman, K.J. et al. Chemical, thermal, and electric field induced unfolding of single protein molecules studied using nanopores. Anal. Chem. 83, 5137–5144 (2011).

36. Arjmandi, N., Van Roy, W., Lagae, L. & Borghs, G. Measuring the Electric Charge and Zeta Potential of Nanometer-Sized Objects Using Pyramidal-Shaped Nanopores. Anal. Chem. 84, 8490–8496 (2012).

37. Pappenberger, G., McCormack, E.A. & Willison, K.R. Quantitative actin folding reactions using yeast CCT purified via an internal tag in the CCT3/gamma subunit. J. Mol. Biol. 360, 484–496 (2006).

38. Muthukumar, M. Theory of capture rate in polymer translocation. J. Chem. Phys. 132 (2010).

39. Waduge, P. et al. Nanopore-Based Measurements of Protein Size, Fluctuations, and Conformational Changes. ACS Nano 11, 5706–5716 (2017).

40. Matsudaira, P., Bordas, J. & Koch, M.H.J. Synchrotron x-ray diffraction studies of actin structure during polymerization. Proc. Natl. Acad. Sci. USA 84, 3151–3155 (1987).

41. Fitzkee, N.C. & Rose, G.D. Reassessing random-coil statistics in unfolded proteins. Proc. Natl. Acad. Sci. USA 101, 12497–12502 (2004).

42. Shortle, D. & Ackerman, M.S. Persistence of native-like topology in a denatured protein in 8 M urea. Science 293, 487–489 (2001).

43. Chou, S.Z. & Pollard, T.D. Mechanism of actin polymerization revealed by cryo-EM structures of actin filaments with three different bound nucleotides. Proc. Natl. Acad. Sci. USA 10, 4265–4274 (2019).

44. Kudryashov, D.S. et al. The crystal structure of a cross-linked actin dimer suggests a detailed molecular interface in F-actin. Proc. Natl. Acad. Sci. USA 102, 13105–13110 (2005).

45. Spector, I., Shochet, N.R., Kashman, Y. & Groweiss, A. Latrunculins: novel marine toxins that disrupt microfilament organization in cultured cells. Science 219, 493–495 (1983).

46. Saito, S.Y. et al. Actin-depolymerizing effect of dimeric macrolides, bistheonellide A and swinholide A. J. Biochem. 123, 571–578 (1998).

47. Stumpe, M.C. & Grubmuller, H. Interaction of urea with amino acids: implications for urea-induced protein denaturation. J. Am. Chem. Soc. 129, 16126–16131 (2007).

48. de Oliveira, G.A.P. & Silva, J.L. A hypothesis to reconcile the physical and chemical unfolding of proteins. Proc. Natl. Acad. Sci. USA 112, E2775–E2784 (2015).

49. Jakova, E. & Lee, J.S. Behavior of alpha-synuclein-drug complexes during nanopore analysis with a superimposed AC field. Electrophoresis 38, 350–360 (2017).

50. Sleutel, M. et al. Nucleation and growth of a bacterial functional amyloid at single-fiber resolution. Nat. Chem. Biol. 13, 902–908 (2017).

51. Olshina, M.A. et al. Plasmodium falciparum coronin organizes arrays of parallel actin filaments potentially guiding directional motility in invasive malaria parasites. Malar. J. 14, 280 (2015).

52. Meller, A. & Branton, D. Single molecule measurements of DNA transport through a nanopore. Electrophoresis 23, 2583–2591 (2002).

