## Supplementary Information for "Single-molecule nanopore sensing of actin dynamics and drug binding"

### Label-free nanopore sensing reveals actin unfolding and real-time drug binding with single-molecule resolution

#### S1. Calibration of the effective length $\boldsymbol{H_{\mathrm{eff}}}$ using 1 kb dsDNA

Unlike solid-state nanopore embedded in a membrane, the effective length ($\boldsymbol{H_{\mathrm{eff}}}$) of nanopipettes is difficult to directly measure or calculate. In addition, since the frequency of the low-pass filter has a significant effect on the amplitudes of current blockade, $\boldsymbol{H_{\mathrm{eff}}}$ of nanopipettes has to be calibrated at the same filter to ensure the calculated excluded volume gives an accurate truth. Calibration using dsDNA molecules has been demonstrated before(1), based on the well-studied excluded atomic volumes of dsDNA. In order to minimise the temporal bias, 1 kb dsDNA was selected to estimate $\boldsymbol{H_{\mathrm{eff}}}$ due to its comparable dwell time (within 0.1 ms) with small proteins and length of 340 nm that exceeds $\boldsymbol{H_{\mathrm{eff}}}$ of nanopipettes. The formula used for calibration was $\Delta \boldsymbol{I_{\mathrm{b}}} \approx \boldsymbol{\sigma\psi} \cdot \boldsymbol{A_{\mathrm{m}}}/\boldsymbol{H_{\mathrm{eff}}}$, where the parameters applied were $\boldsymbol{\sigma} = \mathbf{103.5\ mS/cm}$, $\boldsymbol{\psi} = \mathbf{250\ mV}$, and $\boldsymbol{H_{\mathrm{eff}}} = \mathbf{110\ nm}$ as determined below.

By using the Radical Planes method(2), the atomic volume of AT and GC pairs can be defined as $\boldsymbol{\Lambda}(\mathbf{A+T}) = \mathbf{0.6183\ nm^3}$ and $\boldsymbol{\Lambda}(\mathbf{G+C}) = \mathbf{0.6066\ nm^3}$. The average volume for each base pair $\boldsymbol{\Lambda_{\mathrm{bp}}} = \mathbf{0.61245\ nm^3}$ and the rise in the helix per base pair, 0.34 nm, give the excluded volume per unit length, $\boldsymbol{A_{\mathrm{DNA}}} = \frac{\mathbf{0.61245\ nm^3}}{\mathbf{0.34\ nm}} = \mathbf{1.8\ nm^2}$. This value is in the same dimension as the cross-sectional area based on the crystallography but more applicable to the estimation of excluded volume in nanopore experiments. The $\boldsymbol{H_{\mathrm{eff}}}$ can be determined by the statistical data of peak current in Fig. S9, giving a calibration value of 110 ± 15 nm.

**S2. Calculation of actin unfolding free energy ($\Delta G$)**

First, we determined the free energy at each urea concentration using the equation below:

$$\Delta \boldsymbol{G} = -\boldsymbol{RTln}\frac{[\mathbf{U}]}{[\mathbf{N}]} = -\boldsymbol{RTln}\frac{\boldsymbol{F}_{\mathbf{U}}}{\boldsymbol{F}_{\mathbf{N}}} \quad \text{(S1)}$$

in which $\boldsymbol{F}_{\mathbf{U}}$ and $\boldsymbol{F}_{\mathbf{N}}$ is the fraction of unfolded and native actin, T is given as 298 K. Then by plotting the free energy as a function of urea concentration as shown in Fig. S11, we can obtain the free energy of unfolding at 0 M urea.

$$\Delta \boldsymbol{G} = \Delta \boldsymbol{G}_{\mathbf{U}} - \boldsymbol{m}_{\mathbf{U}}[\mathbf{urea}] \quad \text{(S2)}$$

#### S3. Estimate of the zeta potential for folded and unfolded actin

From protein translocation events, particularly the dwell time, we can estimate the zeta potential of analytes using the equation below:

$$\boldsymbol{\zeta} = \frac{\eta H_{\mathrm{eff}}^2}{\varepsilon} \frac{\partial\left(\frac{1}{t_{\mathrm{d}}}\right)}{\partial \psi} = \frac{\eta H_{\mathrm{eff}}}{\varepsilon} \frac{\partial(v_{\mathrm{eff}})}{\partial \psi} \quad \text{(S3)}$$

where $\boldsymbol{\eta}$ is the solution viscosity, $\boldsymbol{H}_{\mathbf{eff}}$ is the effective length of the nanopore, $\boldsymbol{\varepsilon} = \boldsymbol{\varepsilon}_{\mathbf{0}}\boldsymbol{\varepsilon}_{\mathbf{r}}$ is the solution dielectric constant, $\boldsymbol{t}_{\mathbf{d}}$ is dwell time of protein translocation, $\boldsymbol{\psi}$ is the applied voltage and $\boldsymbol{v}_{\mathbf{eff}}$ is the average effective velocity.

Prior to performing calculation, the physical constants of solutions were calibrated mainly based on the concentration of KCl (3, 4) and urea (5, 6) at 25°C. The viscosity is 0.874 Pa s for 1 M KCl actin monomeric buffer and 1.228 Pa s for the same buffer with 6 M urea. The corresponding dielectric constant ($\boldsymbol{\varepsilon}_{\mathbf{r}}$) is 78.57 and 94.43, respectively.

$\boldsymbol{\partial}(\boldsymbol{v}_{\mathbf{eff}})/\boldsymbol{\partial\psi}$ was extracted from the linear fitting of $\boldsymbol{v}_{\mathbf{eff}}$ vs voltages as shown in Fig. 3b.

**S4. Theoretic models of protein flux in nanopore translocation**

The normalised capture rates ($\boldsymbol{J}_{\mathrm{C}}/\boldsymbol{C}_{\mathbf{0}}$) for protein translocation in nanopore can be generally expressed as

$$\boldsymbol{J}_{\mathrm{C}}/\boldsymbol{C}_{\mathbf{0}} = \frac{\boldsymbol{D}_{\mathbf{m}}}{\boldsymbol{H}_{\mathbf{eff}}}\left[\frac{\boldsymbol{\eta}\left(\boldsymbol{e}^{\widetilde{\boldsymbol{F}}}-\mathbf{1}\right)}{\widetilde{\boldsymbol{F}}} + \frac{(\mathbf{1}-\boldsymbol{\eta})}{(\widetilde{\boldsymbol{F}}+|\tilde{\boldsymbol{z}}\boldsymbol{\psi}|)}\left(\boldsymbol{e}^{\widetilde{\boldsymbol{F}}} - \boldsymbol{e}^{-|\tilde{\boldsymbol{z}}\boldsymbol{\psi}|}\right)\right]^{-\mathbf{1}} \quad \text{(S4)}$$

where $\boldsymbol{C}_{\mathbf{0}}$ is the protein concentration, $\boldsymbol{D}_{\mathbf{m}}$ is the diffusion coefficient of proteins, $\boldsymbol{H}_{\mathbf{eff}}$ is the effective length of nanopores, $\boldsymbol{\eta}$ is the solution viscosity, $\widetilde{\boldsymbol{F}}$ is the total energy barrier, $\tilde{\boldsymbol{z}} = \boldsymbol{\mu}/\boldsymbol{D}_{\mathbf{m}}$ a corrected factor of electrophoretic mobility and $\boldsymbol{\psi}$ is the applied voltages.

**S5. Estimation of protein shapes in nanopore experiments**

Protein hydrodynamic radii ($\boldsymbol{R}_{\mathbf{H}}$), assuming they maintain their globular structures during transport through the nanopore, can be estimated from fractional current amplitude data based on(7, 8)

$$\boldsymbol{R}_{\mathbf{H}} \cong \frac{1}{2}\left[\langle \Delta \boldsymbol{I}/\boldsymbol{I}_{\mathbf{o}}\rangle \left(\boldsymbol{H}_{\mathbf{eff}} + \mathbf{0.8}\boldsymbol{D}_{\mathbf{p}}\right)\boldsymbol{D}_{\mathbf{p}}^{2}\right]^{\mathbf{1/3}} \text{ (S5)}$$

Where $\Delta \boldsymbol{I}$ is the blockade current, $\boldsymbol{I}_{\mathbf{o}}$ is the open current, $\boldsymbol{D}_{\mathbf{p}}$ is the diameter and $\boldsymbol{H}_{\mathbf{eff}}$ is the effective length of a nanopipette.

The theoretical value for $\boldsymbol{\rho}$ is 0.775 for a hard-sphere of uniform density, whereas for an oblate ellipsoid, values of $\boldsymbol{\rho}$ range from 0.875 to 0.987, and for a prolate ellipsoid, values from 1.36 to 2.24.

#### S6. Kinetics measurements of actin-binding with Swinholide A

In a typical kinetic experiment of actin monomer with Swinholide A, the actin monomers in the external bath were induced to translocate through a nanopipette under application of voltages. This procedure lasts at least 1 hr to make sure the protein flux is stable. Without stopping the recording, we then added the Swinholide A to the bath as the time zero point. We counted the dimer number per two minutes with threshold of 75 pA and calculated the population fraction to plot a time-dependent curve. The curves were fitted by a three-component model with positive cooperativity:

$$\boldsymbol{ABC_{\mathrm{t}} = ABC_{\mathrm{max}}\left(1 - e^{-k_{\mathrm{obs}}t}\right)} \text{ (S6)}$$

Where $\boldsymbol{ABC_{\mathrm{max}}}$ is the maximum percentage of the final product, and $\boldsymbol{k_{\mathrm{obs}}}$ is the observed constant.

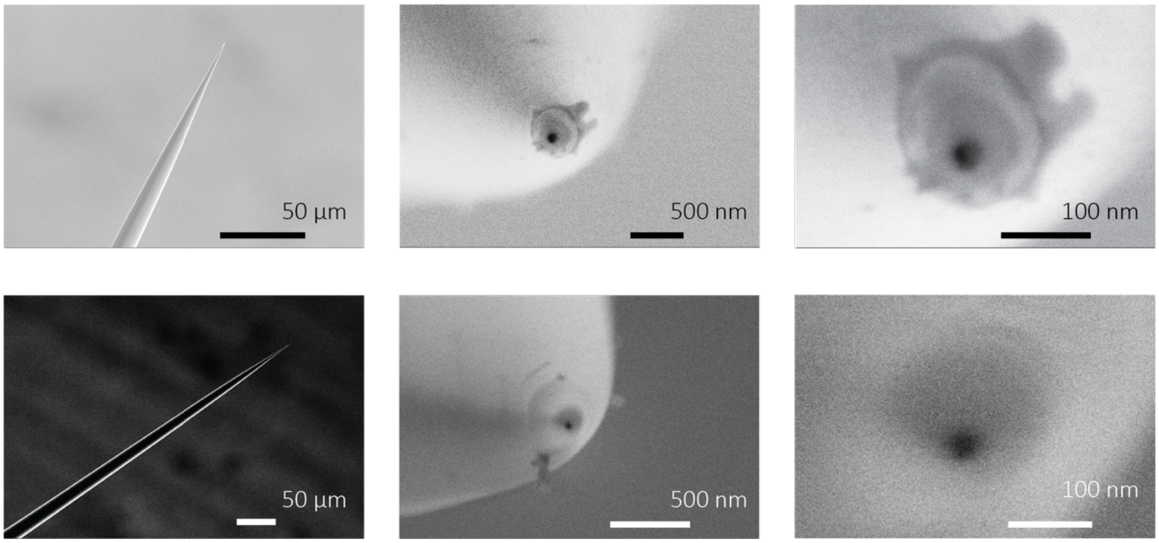


**Fig. S1** SEM images of nanopipettes fabricated by a laser-assisted pulling protocol of quartz capillaries. Left: SEM images showing a conical shape at the tapered tip of nanocapillaries. Middle: deeper landscape of the tip of nanopipettes with a nanoscale pore in the centre. Right: close-up SEM images of the nanopore with a diameter of 25 ± 4 nm formed in the pipettes.

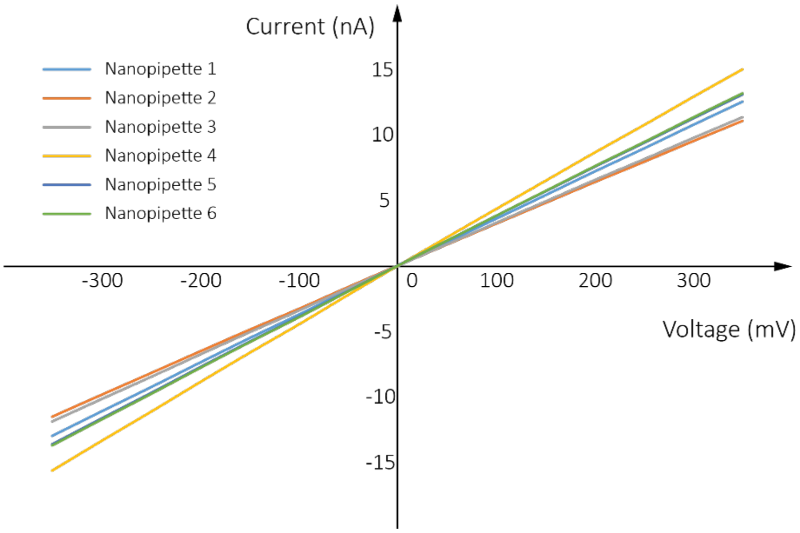


**Fig. S2** Current-voltage curves of nanopipettes used in nanopore measurements. I-V curves obtained in 1 M KCl actin monomeric buffer (containing 20 mM Tris pH 8, 0.4 mM ADP, 0.1 mM $CaCl_2$, 0.01% DMSO and 10% glycerol). The nanopore conductance was estimated to be 37.0 ± 3.9 nS by linear fitting.

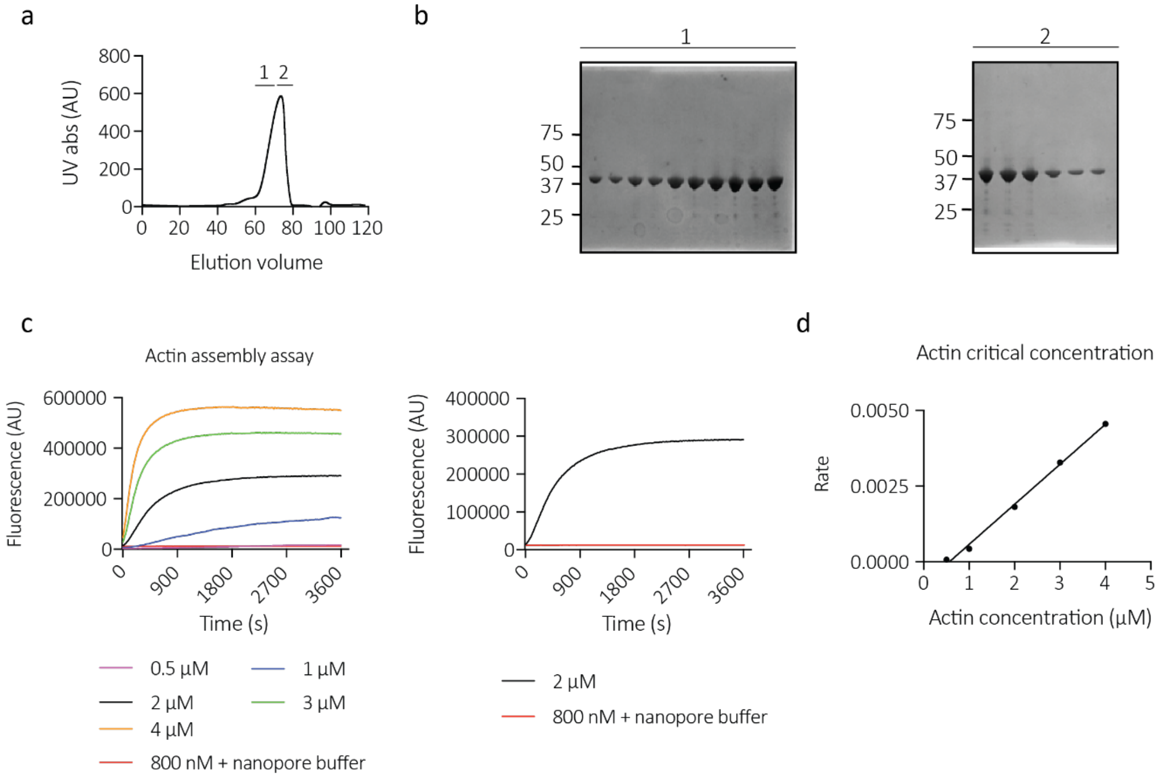


**Fig. S3** Actin was purified as previously described to high purity and was functionally active. **a** A size exclusion chromatography profile of the purified actin. Monomeric actin eluted between elution fractions 60 mL and 80 mL. **b** SDS-PAGE analysis of the monomeric elution fractions. **c** A sample of the actin was labelled with pyrene and checked for activity using a pyrene fluorescence filamentation assay. Actin polymerised as normal under the presence of polymerisation enhancing factors (**c** and **d**) with a critical concentration of 0.6 µM. d and did not form filaments in the nanopore buffer at the concentrations used in the nanopore experiments (**c**).

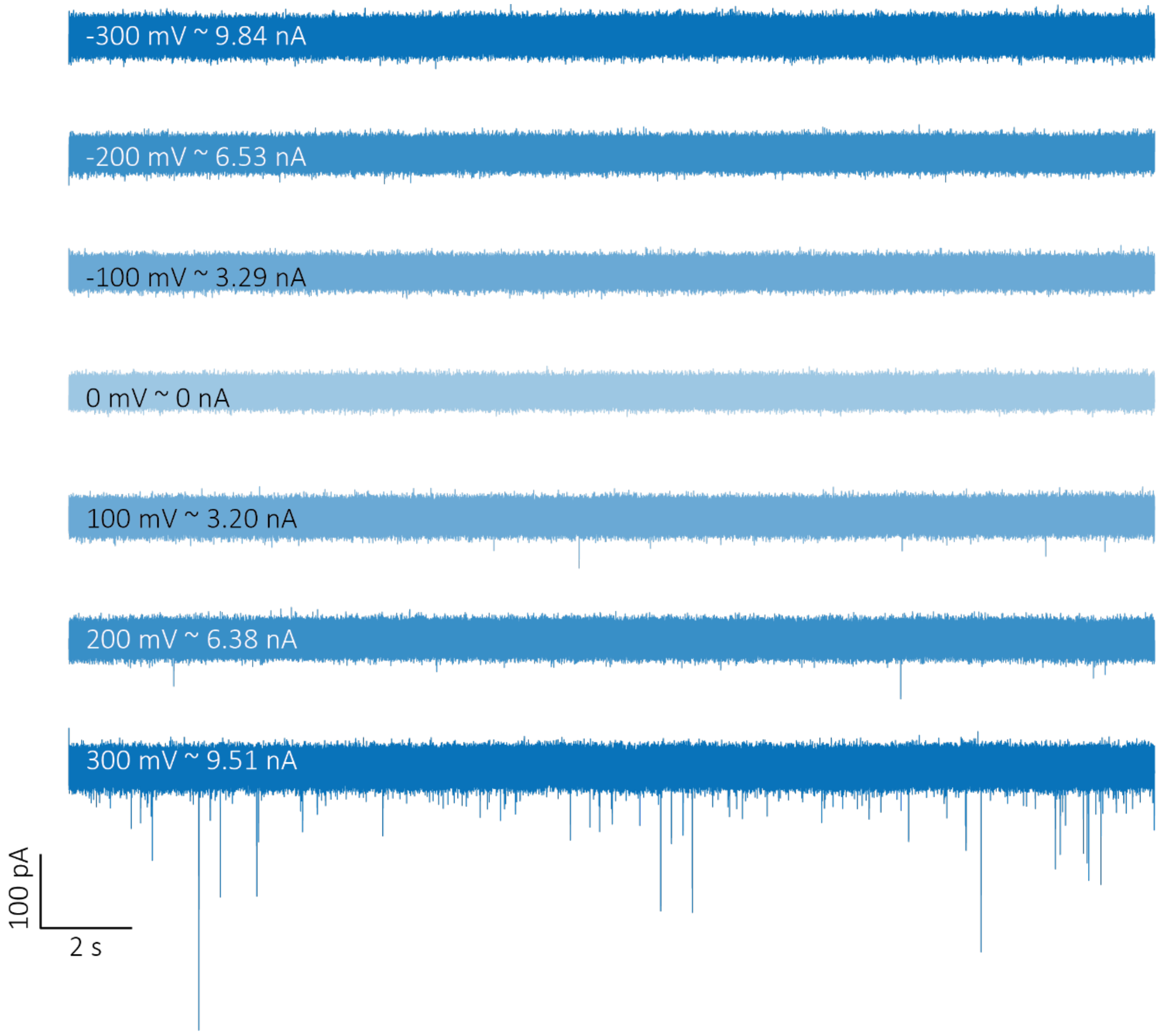


**Fig. S4** Ionic current-time traces of 800 nM actin monomers. The nanopore experiments were conducted in 1 M KCl buffer containing 20 mM Tris pH 8, 0.4 mM ADP, 0.1 mM $CaCl_2$, 0.01% DMSO and 10% glycerol, recorded at 4.17 MHz, filtered to 50 kHz, and collected at different voltages. These traces show translocation spikes at positive voltages with a voltage dependence but no translocation events at negative voltages, demonstrating the EP/EO theory as expected.

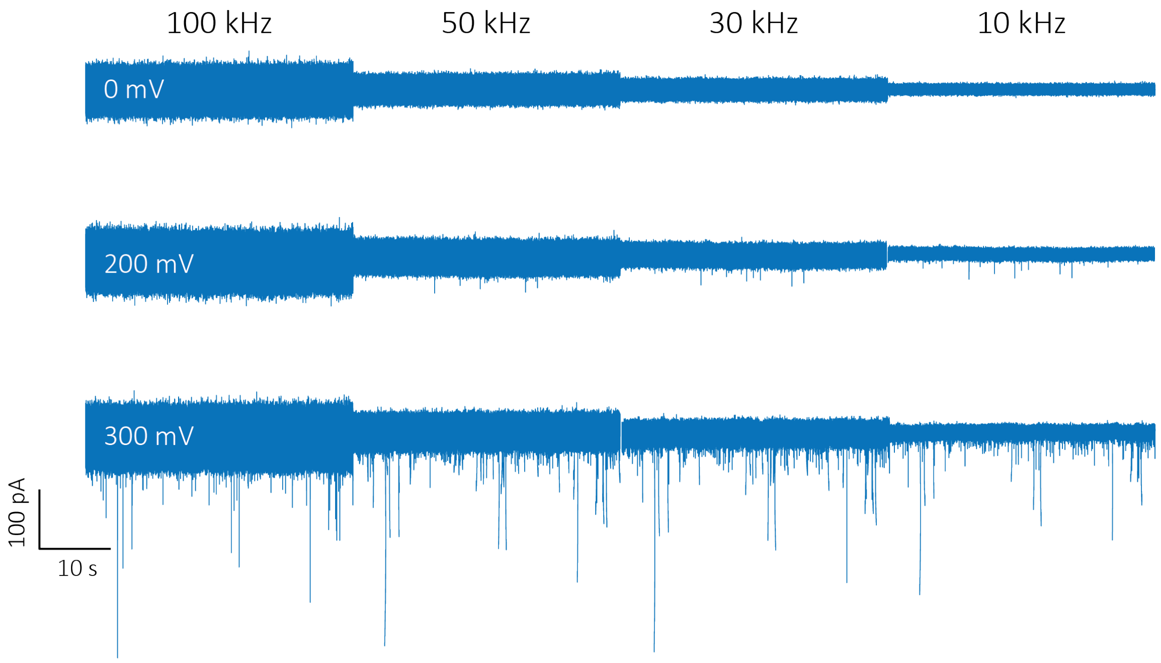


**Fig. S5** Current-time traces showing actin monomers translocation at 0 mV, 200 mV and 300 mV after different low-pass filters. Limited by the signal-to-noise ratio (SNR), no spikes can be observed for 200 mV at 100 kHz filter. Therefore, a low-pass filter of 50 kHz was selected for all data analysis.

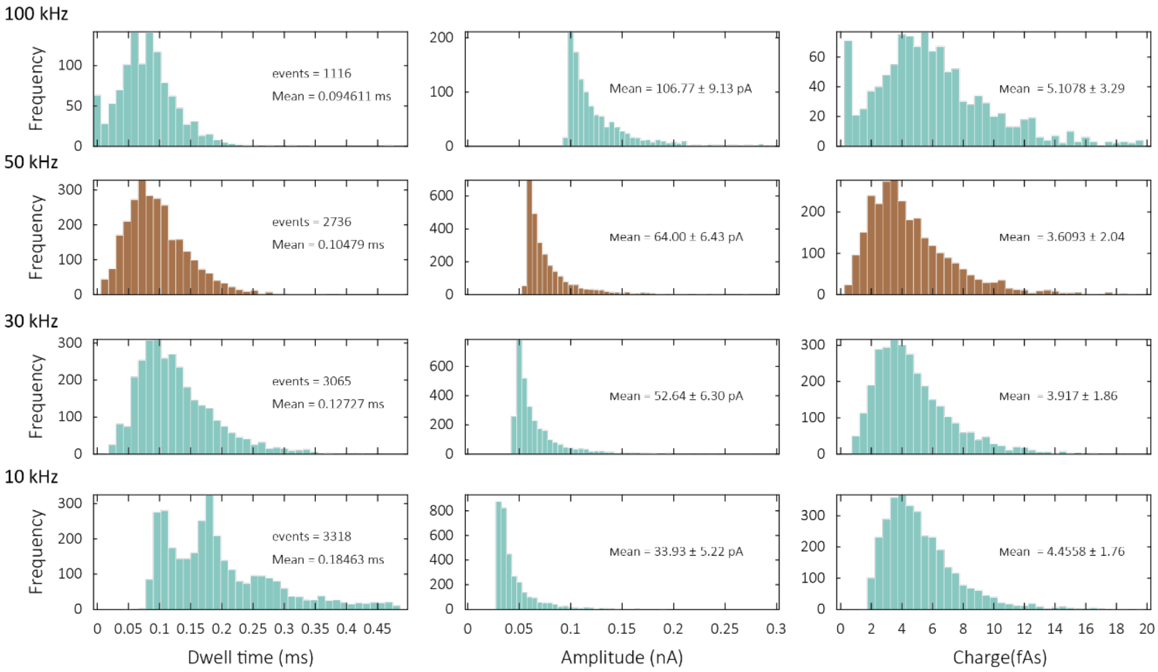


**Fig. S6** Low-pass filter effect on data analysis of 800 nM actin translocation at 250 mV. The frequency of low-pass filter has a significant influence on translocation information, including dwell time, peak current and SNR. At 100 kHz filter, event number is limited by poor SNR and some fake signals with very short dwell time occur due to instrumental noise. At low filter frequency (10 kHz), however, the signals have a bias to real values because of the limitation of bandwidth (sub-population in dwell time histogram). Based on these results, 50 kHz was chosen as the parameter in all data analysis. All statistic in this figure was processed by the same original data.

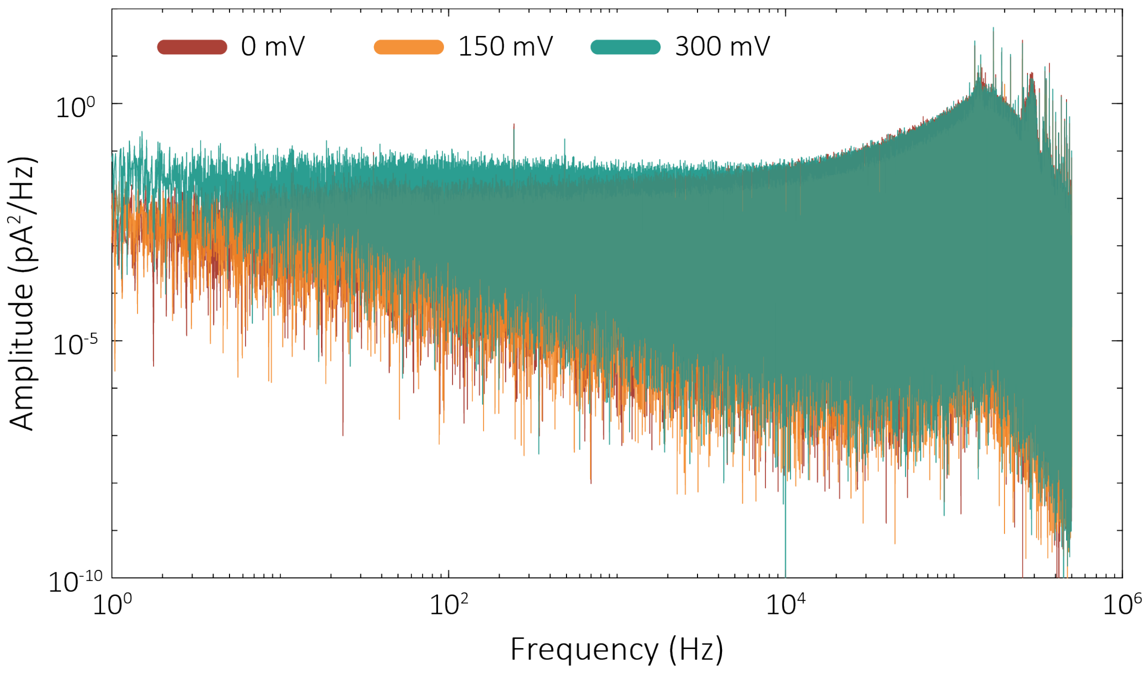


**Fig. S7** Power spectral density (PSD) plots of nanopipette set-up (1 M KCl actin monomeric buffer). PSD plots of nanopipettes at 0, 150 and 300 mV.

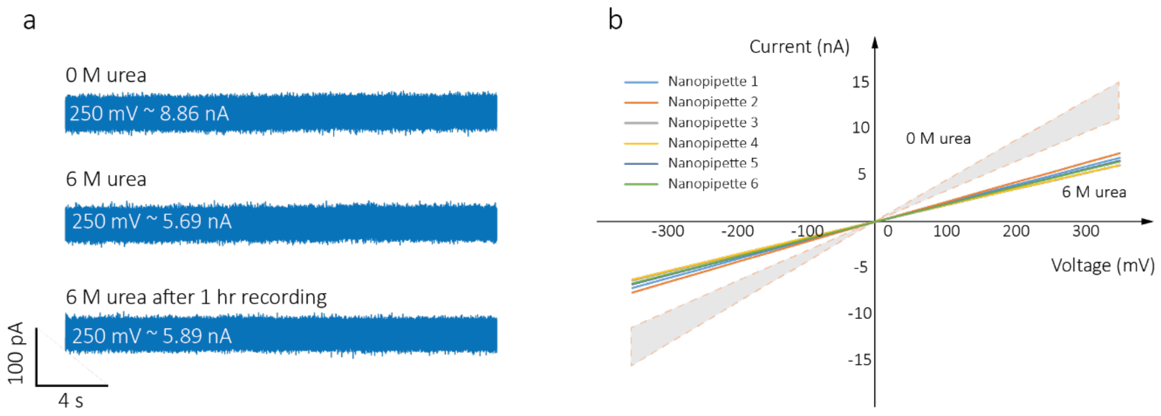


**Fig. S8 a** Current-time traces of 1 M KCl actin monomeric buffer containing 0 M and 6 M urea respectively at 250 mV. The trace for 6 M urea shows a decrease of ionic current due to the decrease of solution conductivity and remains stable after a long-time recording, indicative of a good compatibility of quartz nanopipettes with urea. **b** I-V curves of nanopipettes obtained in 1 M KCl actin monomeric buffer with 6 M urea. The grey shadow is the I-V curves tested in 0 M buffer.

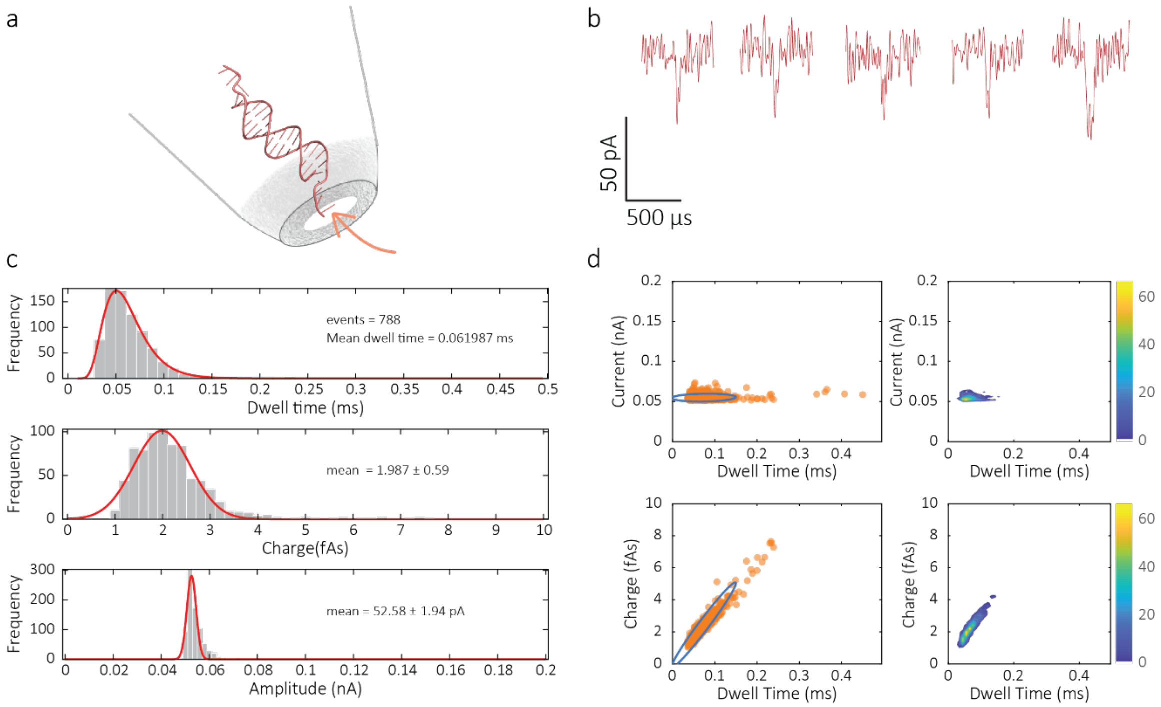


**Fig. S9** 100 pM 1 kb dsDNA translocation statistic at 250 mV. This experiment was conducted in the same buffer of actin monomers (1 M KCl) and the same out-to-in mode. **a** Nanopore experimental configuration of 1 kb DNA translocation. **b** Typical translocation events at a low-pass filter of 50 kHz. **c** Histograms of dwell time, charge and peak current distribution for 1 kb dsDNA translocation at 250 mV. **d** Scatterplots and 2D-Kernel plots of dwell time vs. peak current and charge for 1 kb dsDNA.

| | Conductivity (mS/cm) |
|---|---|
| Buffer* | 103.5 |
| Buffer + 1 M urea | 101.9 |
| Buffer + 2 M urea | 96.4 |
| Buffer + 3 M urea | 85.1 |
| Buffer + 4 M urea | 74.9 |
| Buffer + 5 M urea | 68.8 |
| Buffer + 6 M urea | 65.8 |

Buffer* contains 1 M KCl, 20 mM Tris pH 8, 0.4 mM ADP, 0.1 mM $CaCl_2$, 0.01% DMSO and 10% glycerol.

**Table S1** Solution conductivity for buffers with different concentrations of urea used in actin unfolding assays.

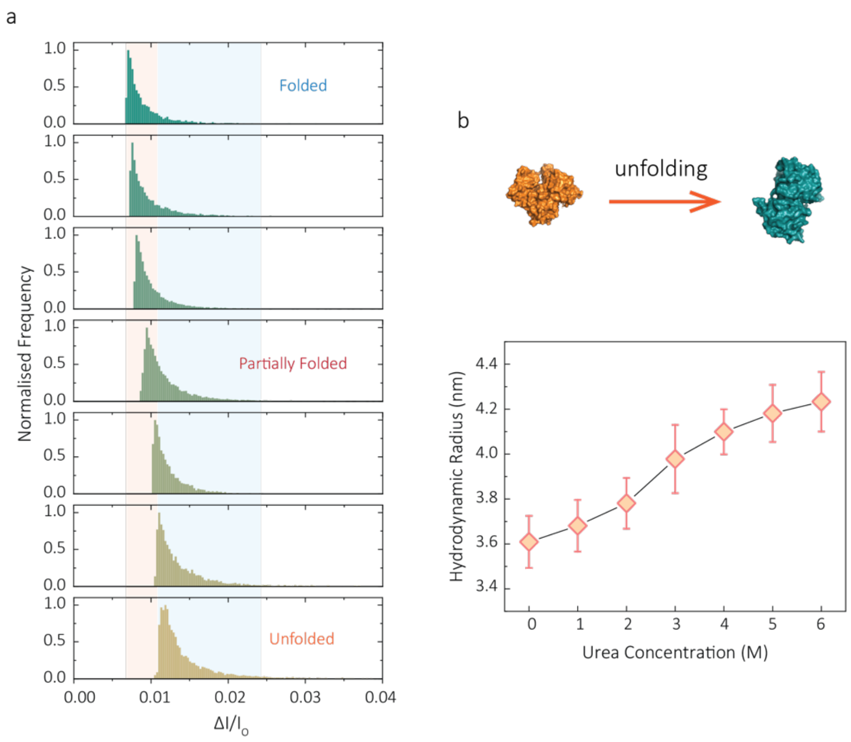


**Fig. S10** Changes of hydrodynamic radii ($\boldsymbol{R}_\mathbf{H}$) of actin monomers during unfolding process. **a** Normalised distribution of fractional current blockade ($\Delta\boldsymbol{I}/\boldsymbol{I}_\mathbf{o}$) in different urea concentrations. The orange boundary represents the fully folded actin state, and blue boundary determines the unfolded actin or other transient aggregates at low urea concentrations. The population shifts from a mostly native, folded state to a higher excluded volume, consistent with an unfolded state with larger hydrodynamic radii, as the urea concentration increases. **b** Top: schematic of conformational changes during actin unfolding. Bottom: plot of $\boldsymbol{R}_\mathbf{H}$ vs urea concentrations showing a two-state trend.

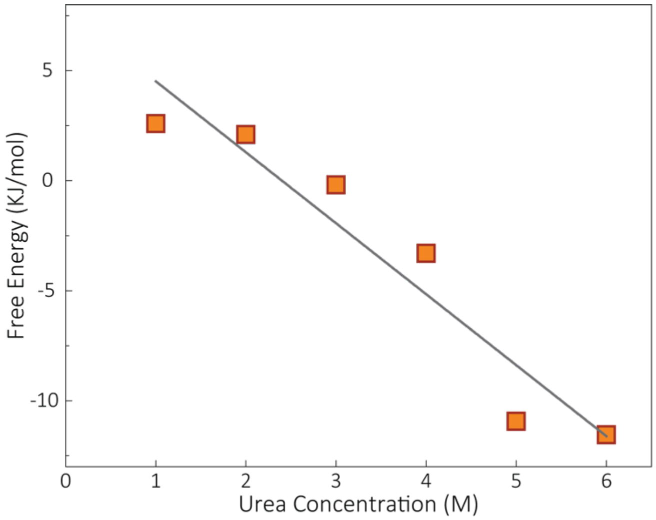


**Fig. S11** Plots of free energy as a function of urea concentration along with a linear fitting.

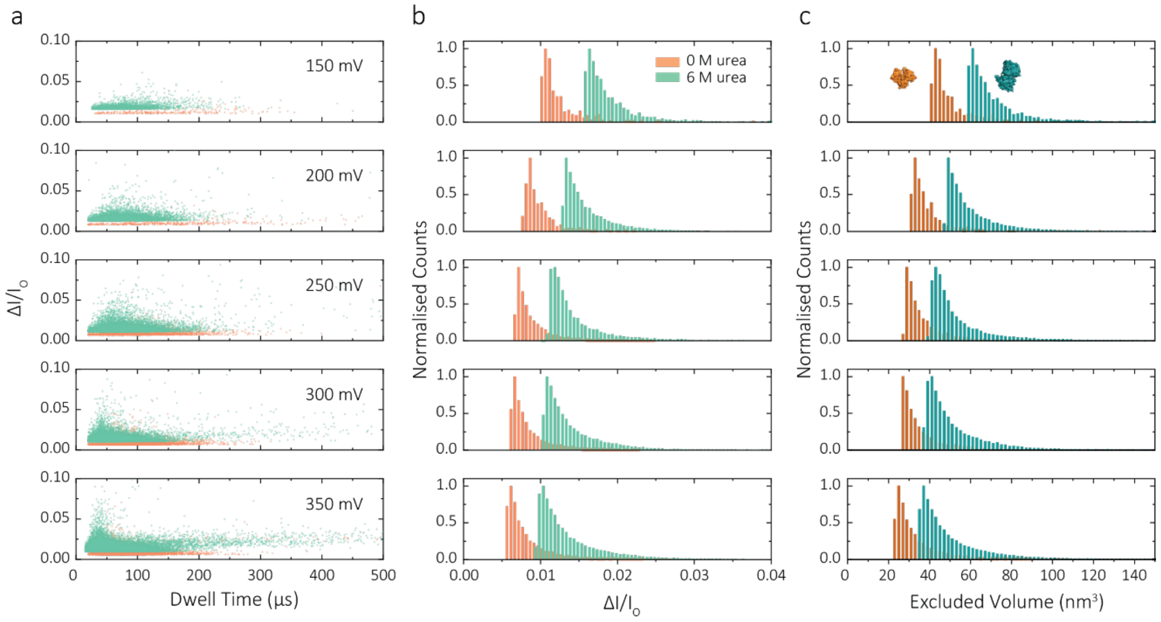


**Fig. S12** Discriminating folded and unfolded actin using nanopipettes performed in 1 M KCl with 0 M or 6 M urea. **a** Scatterplots of fractional current blockades vs dwell times for folded and unfolded actin at different voltages. **b** Distributions of fractional current blockades for both folded and unfolded actin at various voltages. **c** Statistic histograms of the excluded volume calculated from eq. 1 for folded and unfolded actin at different voltages from 150 mV to 350 mV.

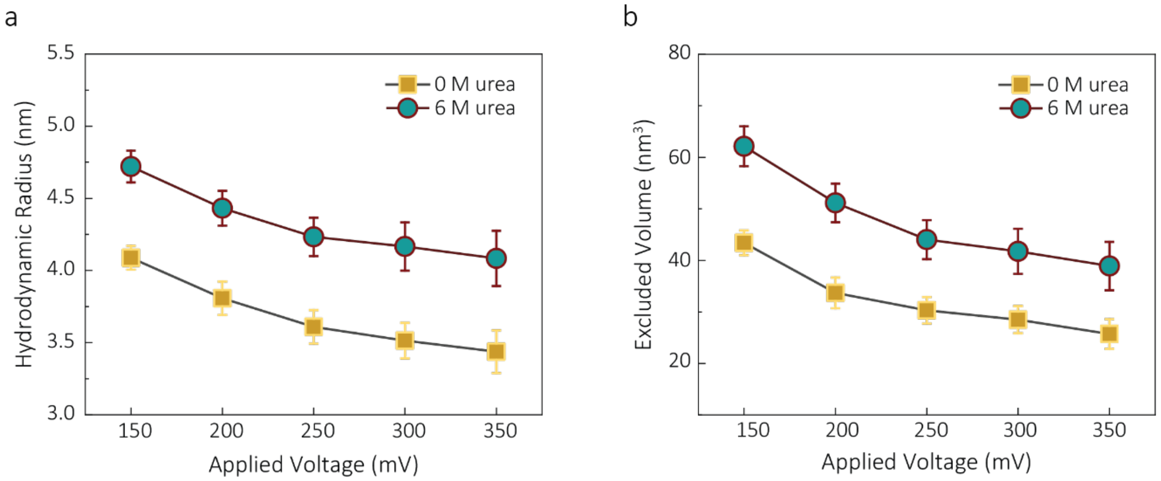


**Fig. S13 a** Plots of hydrodynamic radius and **b** excluded volume vs voltages for folded and unfolded actin.

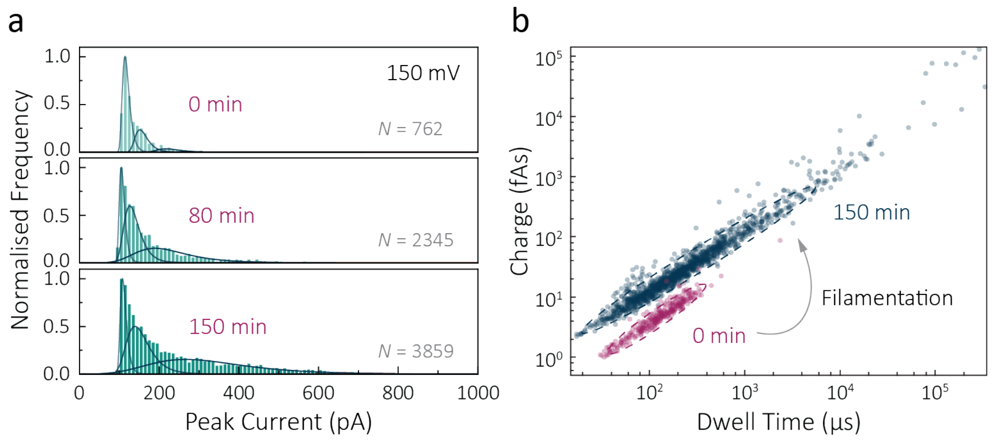


**Fig. S14 a** Histograms of the peak current for actin filament formation at 0, 80 and 150 min at 150 mV, showing a distribution of multiple populations and an extension in higher peak currents. **b** Scatterplots of charge vs dwell time for actin polymerisation at 0 and 150 min.

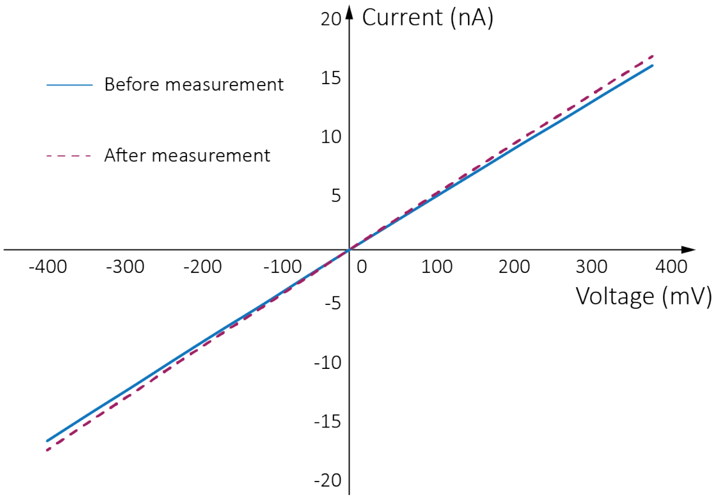


**Fig. S15** IV curves before and after measurement of actin filament formation (1 µM monomer concentration in ATP buffer). The slight increase of conductivity is indicative of water evaporation over time and no interaction between protein and nanopore surface.

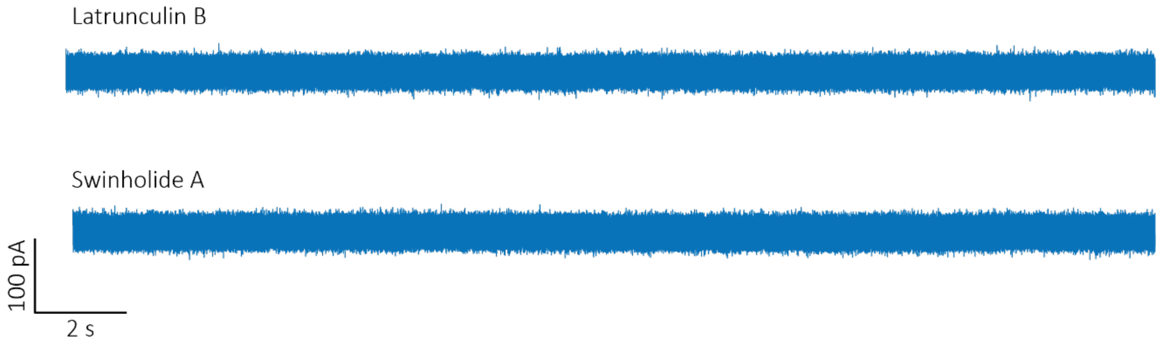


**Fig. S16** Latrunculin B and Swinholide A do not interact with the nanopipette surface. Current-time trances show no abnormal noise for both drugs at 250 mV.

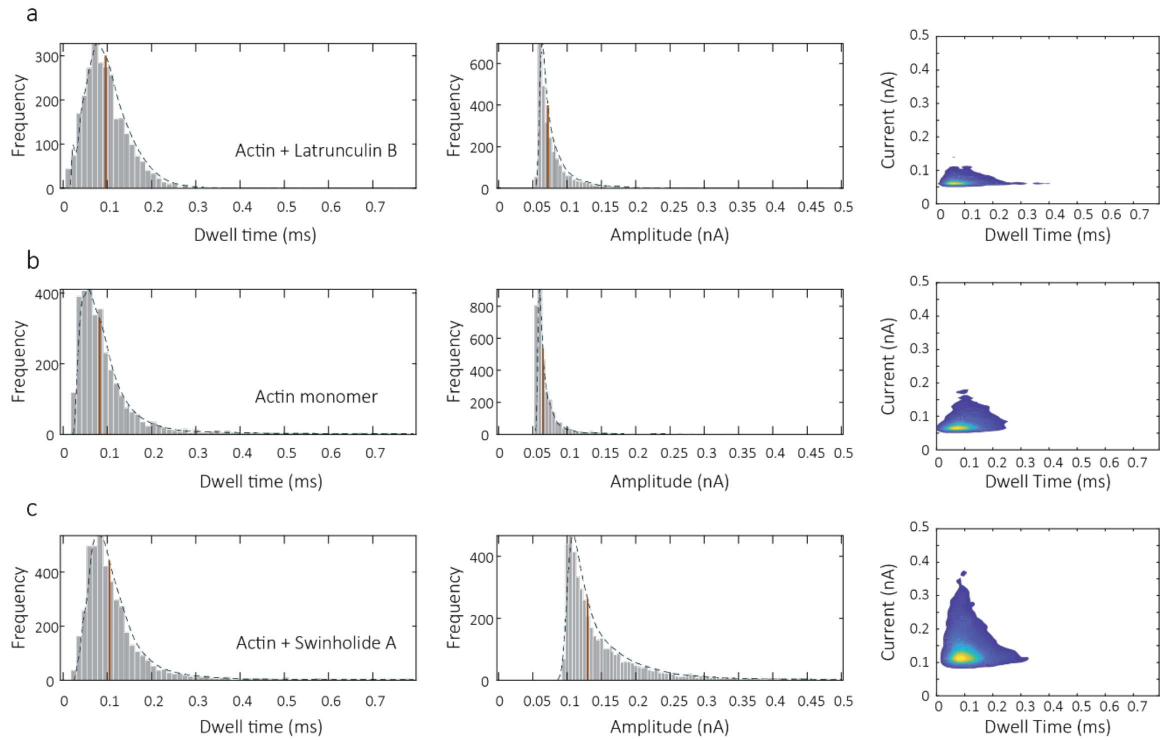


**Fig. S17** Statistic information for **a** Actin bound with Latrunculin B, **b** Actin monomers and **c** Actin bound with Swinholide A. Left: distribution of dwell time. Middle: distribution of peak current. Right: 2-D Kernel plots of peak current vs dwell time. All data were collected in 1 μM actin and 10 μM drug at 250 mV.

#### References


1. D. S. Talaga, J. Li, Single-molecule protein unfolding in solid state nanopores. *J. Am. Chem. Soc.* **131**, 9287-9297 (2009).
2. K. Nadassy, I. Tomas-Oliveira, I. Alberts, J. Janin, S. J. Wodak, Standard atomic volumes in double-stranded DNA and packing in protein-DNA interfaces. *Nucleic Acids Res.* **29**, 3362-3376 (2001).
3. D. E. Goldsack, A. A. Franchetto, The viscosity of concentrated electrolyte solutions—III. A mixture law. *Electrochim. Acta* **22**, 1287-1294 (1977).
4. F. H. Drake, G. W. Pierce, M. T. Dow, Measurement of the Dielectric Constant and Index of Refraction of Water and Aqueous Solutions of KCl at High Frequencies. *Phys. Rev.* **35**, 613-622 (1930).
5. K. Kawahara, C. Tanford, Viscosity and density of aqueous solutions of urea and guanidine hydrochloride. *J. Biol. Chem.* **241**, 3228-3232 (1966).
6. J. Wyman, Dielectric Constants: Ethanol—Diethyl Ether and Urea—Water Solutions between 0 and 50°. *J. Am. Chem. Soc.* **55**, 4116-4121 (1933).
7. J. Larkin, R. Y. Henley, M. Muthukumar, J. K. Rosenstein, M. Wanunu, High-bandwidth protein analysis using solid-state nanopores. *Biophys. J.* **106**, 696-704 (2014).
8. P. Waduge *et al.*, Nanopore-Based Measurements of Protein Size, Fluctuations, and Conformational Changes. *ACS Nano* **11**, 5706-5716 (2017).
